# The competition dynamics of resistant and sensitive infections

**DOI:** 10.1101/2021.01.25.427822

**Authors:** T. E. Lee, S. Bonhoeffer, M. A. Penny

## Abstract

Antimicrobial resistance is a major health problem with complex dynamics. Resistance may occur in an area because treated infections mutated and developed resistance, and the proportion of infections in a population may then increase. We developed a novel and flexible model that captures several features of resistance dynamics and competition. The model is able to account for many antimicrobials and thus can generally explore competition dynamics and their impact on pathogens and bacteria.

Unlike simpler models, our nested model allows the population of resistant pathogen to smoothly increase or decrease. Time dependent dynamics are incorporated into difference equations which examines the effects of 12 parameters. This enables us to explicitly include three key competition dynamics: the transmission cost of resistance that occurs between hosts, the fitness cost of resistance that occurs within untreated hosts, and the release of this competition (from the fitness cost) that occurs once a host is treated. For malaria, our results suggest that without competitive release, drug resistance does not emerge. However, once emerged, competitive release has little effect, and the best way to mitigate the spread is to ensure that treatment is very effective.

## Introduction

Antimicrobial resistance (AMR) is the ability of a microorganism (such as bacteria, viruses, and some parasites) to reduce the effect of an antimicrobial (such as antibiotics, antivirals and antimalarials) on the microorganism [1]. The frequency of resistance in a region, and whether presence is due to mutation or transmission, can be informed by data [2]. However, this distinction can be difficult without reasonable genetic data and longitudinal follow up of treated patients.

More generally, the dynamics that contribute to the spread, or inhibition of resistance, and the interplay between these dynamics, are not well understood. Modelling provides a framework to explore different drivers of resistance emergence and its onward transmission. However, many models either ignore co-existence of different genotypes within hosts, or include it but biases in favour of resistant genotypes are introduced. Moreover, often the models are susceptible-infected-susceptible (SIS) compartmental models, and thus ignore the effect of treatment. See [3] for a review on infection models in evolutionary epidemiology.

Here we present a general model of drug sensitive and drug resistant genotypes which incorporates time dependent processes into a susceptible-infected-treated-susceptible (SITS) model. The model has a unique balance of complexity which captures dynamics that affect the sensitive and resistant genotypes within host and between host, and tractability which provides new insight into the influence of different processes on the establishment and spread of resistant genotypes. In particular, the model focuses on the effect, and interplay of three competition dynamics: transmission cost, fitness cost, and competitive release. Transmission and fitness costs are a disadvantage to resistant genotypes, whereas competitive release is an advantage. These dynamics are discussed in more detail in the Methods section.

Due to the novelty of our model, it is particularly relevant to describe how our model relates to current mixed infection models. This exercise explains not only the motivations for our model, but also the specific assumptions and details. Therefore we now provide a brief overview of mixed infection models, and then we describe two major motifs of our nested, co-infection model.

When studying the evolution of a pathogen, there are within host and between host effects that occur on different time scales. Nested models are useful to collate these effects. The importance of nested models to improve our understanding of pathogen evolution was studied in a review [4]. This review concludes that nested models should include a reciprocal feedback between within host dynamics and between host dynamics. This may initially be an obvious statement, however many nested models [5–14] simply use the within host dynamics to inform the between host dynamics, but not vice-versa. A straightforward example is when the model only uses the pathogen load within hosts to inform the between host dynamics, such as in [11, 12, 15, 16]. In these models the transmission probability is directly related to the pathogen load. This relationship is recognised in the transmission of malaria and dengue from humans to mosquitoes [17, 18], the vertical transmission of hepatitis B virus between mothers and infants [19], and the transmission of *Escherichia coli* in cattle [20]. Moreover, pathogen load is generally relatively easy to measure. However, whilst only using pathogen load is intuitively appealing, it is likely to be a poor predictor of infectiousness when used alone [21]. Symptoms which increase transmission are not always directly related to pathogen load, such as ulcers of an HIV-infected host [22]. Moreover, continuing the example of HIV, host-behaviour may decrease transmission via the use of condoms. We note that our model includes reciprocal feedback which is independent of pathogen load.

Sensitive and resistant genotypes coexist at the population level, yet models struggle to reproduce this observation [23]. Mixed infection models are either super-infection models, where one infection replaces the other [24, 25], or co-infection, where hosts may carry sensitive and resistant pathogens simultaneously. Super-infection models consider competition between the two strains or genotypes at the population level, whereas co-infection models consider competition within host and between host. Models may refer to strains or genotypes with different virulence and transmission ability [26], and thus treatment is excluded. We briefly describe how two key super-infection models obtained coexistence at the population level by considering within host dynamics, and then describe co-infection models and challenges that they incur. Our model is a co-infection model, however we also include within host dynamics, similar to the super-infection model which we now describe.

Super-infection models which track resistance have a minimum of of five host compartments: susceptible, infected with the sensitive genotype, treated - carrying sensitive genotype, infected with the resistant genotype, treated - carrying resistant genotype. The number of compartments increases if immune hosts and/or partially resistant genotypes are included. It can be challenging to develop a model which predicts sensitive and resistant genotypes coexisting in the population. An early model [48] accounts for two within-host processes: to capture the fitness cost of resistant bacteria, within untreated hosts the frequency of resistant bacteria is in a state of steady decline. And to capture the release from this competition, once treated, the model assumes that the frequency of resistant bacteria in a treated host goes to 100%.

Whilst this model obtains stable coexistence at the population level, co-infection [5, 28] within hosts is ignored. There is clearly a tractable advantage with modelling super-infection, and not co-infection, especially as the number of possible infections within a host increases. For example, in high transmission settings of malaria, it is common for hosts to carry up to ten infections simultaneously [29]. Modelling this setting requires 125 compartments (excluding compartments for recovered hosts, and partially-resistant infections), which further explodes if the order which hosts are infected by different clones is important. This computational drawback of a compartmental model can be overcome by using an agent based model. However, agent based models for infectious diseases usually include tens of parameters, making computation slow, and thus inhibiting exploration of the parameter space. Moreover, the added complexity requires many assumptions on the dynamics, which may also introduce biases.

Modelling two infections simultaneously is manageable, and thus common. The simplest co-infection models have host compartments: susceptible, infected with sensitive genotype {*S*}, infected with resistant genotype {*R*}, and infected with both {*SR*}. Therefore these models assume that co-infection of the same genotypes do not occur. The review paper [3] demonstrates, using a neutral mutant genotype, that not accounting for sensitive genotypes co-infecting a host unrealistically gives mutant genotypes a fitness advantage. Despite this bias, models of this form are common, for example [28, 30–32]. The first model to correct this bias is [5]. The authors assume co-infection of sensitive genotypes {*SS*}, but not resistant genotypes {*RR*} as they assume co-infection of two rare genotypes is negligible. However, [33] argues that in the case of antibiotics at least, that ignoring co-infection of the same strains, {*SS*} and {*RR*}, implicitly makes ecological assumptions which do not seem biological plausible. In addition, [5] is one of the nested models mentioned previously that does not use the between host dynamics to inform the within host dynamics [4].

We now describe two major motifs of our new model. The first pertains to achieving a high-level overview, but with reduced dependence on unknown assumptions. That is, to avoid modelling processes that would require a model in itself, we use simple variable functions to represent complex processes. Therefore we capture the relevant properties which would emerge from several additional models, and thus we can include many processes in one model in a tractable manner. A major example is that we do not explicitly model the within host processes. This makes the model flexible enough to capture any infection since infection dependent processes such as cells becoming infected, and the consequential immune responses, are not explicitly modelled. And in fact, within host models that capture these processes have had little impact since there remain many forces that are unknown [34]. As such we focus on the main effects that alter the proportion of resistant pathogen, such as the competition dynamics, and omit immune responses, as in [13, 35]. The difference in fitness between the sensitive and resistant genotypes is a simple function inspired by plots of the pathogen load in mice infected with both sensitive and resistant malaria infections [38], such that we allow the fitness cost of resistant genotypes to vary between the extremes of non-existent (‘turned off’) to a fitness cost that is so great that resistant genotypes are unable to survive when sensitive genoytypes are also within an untreated host. Like [48], we do not explicitly model the pharmacokinetics/pharmacodynamics (PK/PD) dynamics. However, whereas [48] assume that after treatment resistant bacteria within hosts simply increases to 100% due to the release of competition with sensitive bacteria, we consider an incremental daily increase which depends on the drug concentration within the host, which changes daily according to the properties of the drug such as its half-life.

Thus competitive release is an emergent property that we have can alter, or again, completely ‘turn off’. Lastly, an example particularly relevant to malaria, we do not explicitly model recombination, a process that occurs within infected mosquitoes (described more in the Methods section). Recombination is relevant because it affects the resistance of the infection that is transmitted. We model this transmission cost as simply a filtration process which affects the probability of resistant pathogen being transmitted from human to human. Again, this makes the model easily adaptable to other diseases whose resistant pathogen population is changed during transmission. Or, again, it may be ‘turned off’, if the infection incurs no transmission cost.

The second motif is, as previously mentioned, our model is independent of pathogen load. Instead we model the proportion of the pathogen population (PRP) which is resistant. By modelling the proportion of the pathogen population which is resistant, within hosts and at the population level, as a continuous variable, the number of infections that a host can carry is not limited. This is especially useful when modelling infections where hosts carry more than two infections simultaneously, such as malaria [29]. Moreover, the competition dynamics between sensitive and resistant strains can be modelled as a continuous process, which is often more suitable than the two-class approach of super-infection models {*S, R*}, or the three-class approach of most co-infection models {*S, SR, R*}. For example, in a mice study [38], two malaria infections within a mouse compete, such that the proportion of parasitaemia which is resistant changes continuously in time, so a discretisation of only two or three classes is coarse. Over the infection time, the mixed parasitaemia density within the mouse is unlikely to be exactly 50/50. Notwithstanding, our model can be parameterised in a manner that mimics a co-infection model with only three classes, {*SS, SR, RR*}. Note that interpreting our model in this manner requires the assumption that hosts carry two infections exactly. Whilst this is a caveat, omitting the possibility of carrying two infections of the same class is also biased [3, 23].

Our nested model, with flexible components, represents many mechanisms of pathogen dynamics, see Fig 1. We run 3000 simulations of the model with varying input values, see Table 1. The output of interest is the PRP, at the population level, at 20 years. By conducting a sensitivity analysis we identify factors and dynamics that determine

- resistance establishment: whether the population PRP is above zero,
- resistance spread: given establishment, what value is the population PRP.

**Figure 1.**
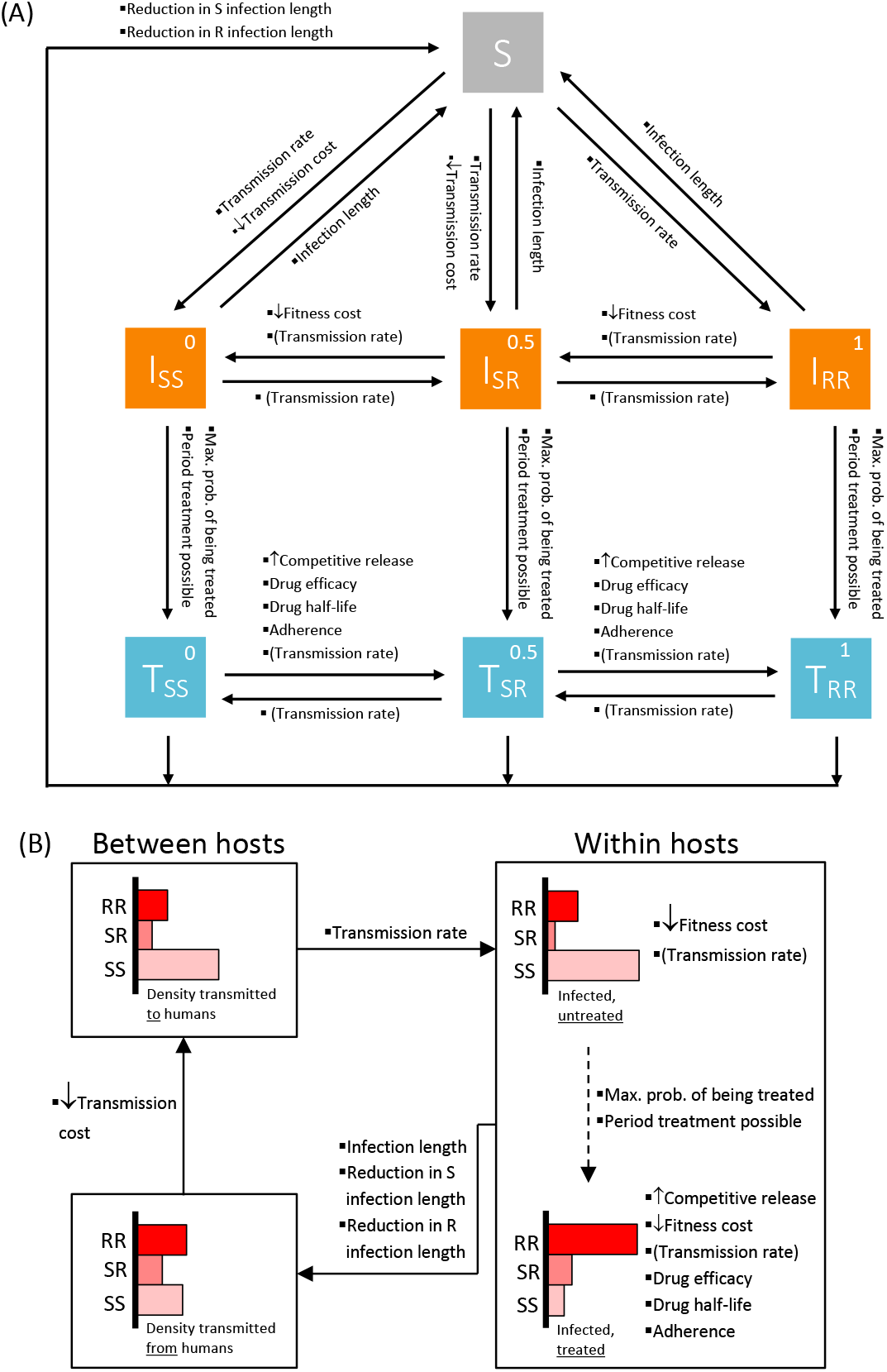
Illustrative model schematics showing (A) how one humans moves through the compartments, and (B) how the overall proportion of resistant pathogens (PRP) in the population are altered throughout the transmission cycle. For clarity, the schematics refer to the case where the PRP has only three levels (from *N* = 2) which correspond to only resistant pathogen, 1:- {*RR*}, half resistant pathogen, 0.5:- {*SR*}, and only sensitive pathogen, 0:- {*SS*}. However, throughout the paper we take *N* = 10. Arrows represent movement through the transmission cycle, and rather than rates, listed besides the arrows are the factors/dynamics that influence this movement. The text-arrows indicate the influence on increasing (↑) or decreasing (↓) resistance in the population. The bracketed ‘(Transmission rate)’ represents when the Transmission Rate has an influence because of multiple transmission events (someone infected can receive further infections). In (A), the PRP within a human is given in the top right of the square for the state (S:- susceptible, I:- infected, T:- treated). In (B), at the population level, the PRP is shown as a sideways histogram. The factors/dynamics listed besides histograms indicate that they impact the PRP during this stage. The dashed arrow indicates that not all infected humans will be treated. On a given day, the distribution that is transmitted to newly infected humans is the same.

**Table 1.**
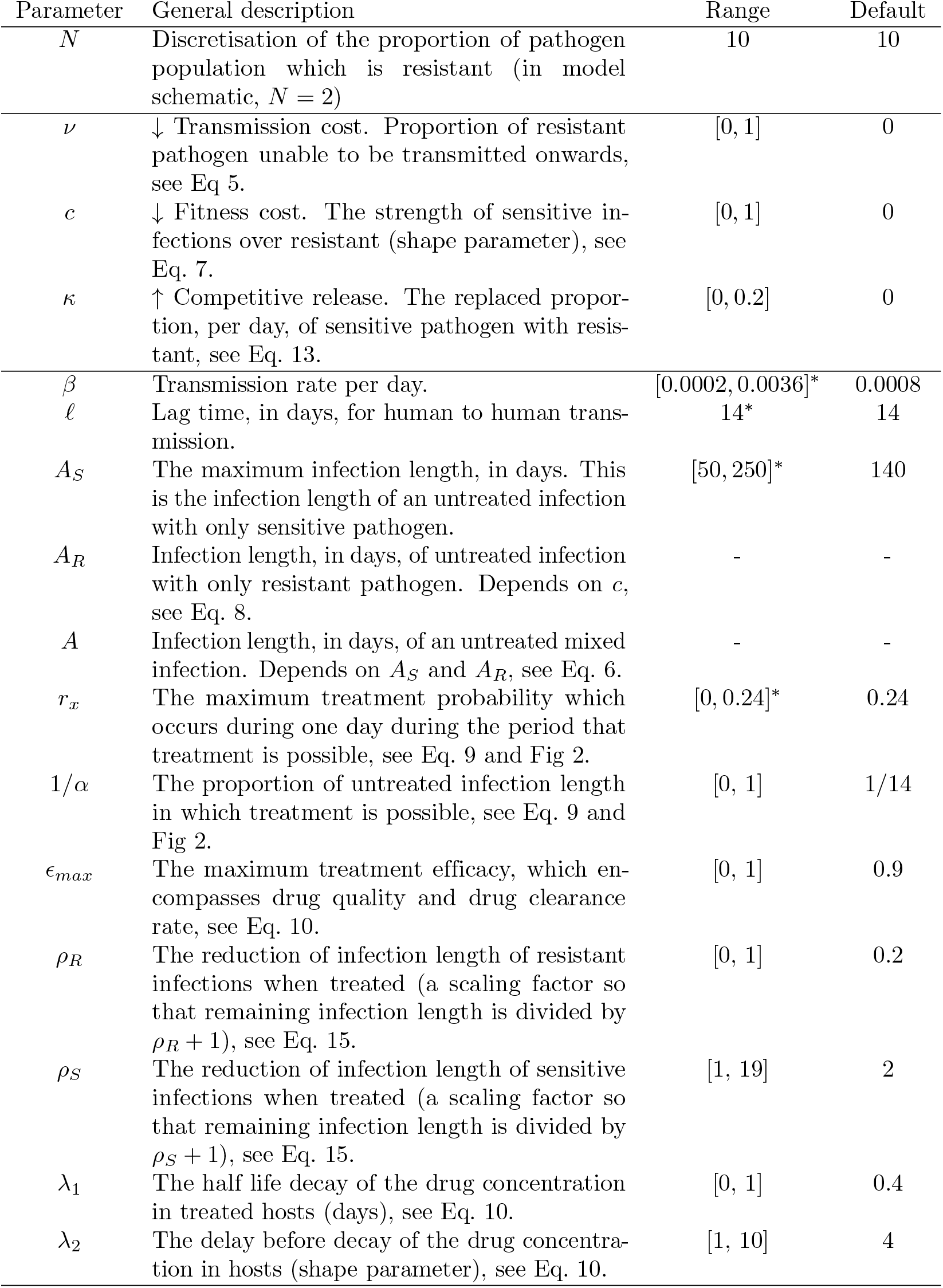
A description of the model parameters, the range chosen for our simulations, and the default value. The asterisk indicates when the ranges are chosen according to malaria parameters. Otherwise, the values are general. The infection lengths *A_R_* and *A* differ for each simulation, but they depend on other parameters so their range is not given in this table.

| Parameter | General description | Range | Default |
| --- | --- | --- | --- |
| $N$ | Discretisation of the proportion of pathogen population which is resistant (in model schematic, $N = 2$) | 10 | 10 |
| $\nu$ | ↓ Transmission cost. Proportion of resistant pathogen unable to be transmitted onwards, see Eq 5. | $[0, 1]$ | 0 |
| $c$ | ↓ Fitness cost. The strength of sensitive infections over resistant (shape parameter), see Eq. 7. | $[0, 1]$ | 0 |
| $\kappa$ | ↑ Competitive release. The replaced proportion, per day, of sensitive pathogen with resistant, see Eq. 13. | $[0, 0.2]$ | 0 |
| $\beta$ | Transmission rate per day. | $[0.0002, 0.0036]^*$ | 0.0008 |
| $\ell$ | Lag time, in days, for human to human transmission. | 14* | 14 |
| $A_S$ | The maximum infection length, in days. This is the infection length of an untreated infection with only sensitive pathogen. | $[50, 250]^*$ | 140 |
| $A_R$ | Infection length, in days, of untreated infection with only resistant pathogen. Depends on $c$, see Eq. 8. | - | - |
| $A$ | Infection length, in days, of an untreated mixed infection. Depends on $A_S$ and $A_R$, see Eq. 6. | - | - |
| $r_x$ | The maximum treatment probability which occurs during one day during the period that treatment is possible, see Eq. 9 and Fig 2. | $[0, 0.24]^*$ | 0.24 |
| $1/\alpha$ | The proportion of untreated infection length in which treatment is possible, see Eq. 9 and Fig 2. | $[0, 1]$ | 1/14 |
| $\epsilon_{max}$ | The maximum treatment efficacy, which encompasses drug quality and drug clearance rate, see Eq. 10. | $[0, 1]$ | 0.9 |
| $\rho_R$ | The reduction of infection length of resistant infections when treated (a scaling factor so that remaining infection length is divided by $\rho_R + 1$), see Eq. 15. | $[0, 1]$ | 0.2 |
| $\rho_S$ | The reduction of infection length of sensitive infections when treated (a scaling factor so that remaining infection length is divided by $\rho_S + 1$), see Eq. 15. | $[1, 19]$ | 2 |
| $\lambda_1$ | The half life decay of the drug concentration in treated hosts (days), see Eq. 10. | $[0, 1]$ | 0.4 |
| $\lambda_2$ | The delay before decay of the drug concentration in hosts (shape parameter), see Eq. 10. | $[1, 10]$ | 4 |

We also investigate different settings, such as low to high treatment, and low to high transmission. From our model we learn about misconceptions that may arise when models exclude certain competition dynamics.

We parameterised the model to malaria since this model is particularly suited to an infection with complex aspects to its transmission, including recombination. Moreover, resistance to artemisinin and several partner drugs is of increasing concern. Nonetheless, the model is readily applicable to strains of other microbials, see Table 1 for more details.

## Methods

We developed a susceptible-infected-treated-susceptible (SITS) compartmental model with additional time dependent dynamics to capture how the mix of sensitive and resistant genotypes vary after a transmission event, and after treatment. The proportion of humans in each compartment, susceptible, infected and treated, is denoted by *S*(*t*), *I*(*t*) and *T* (*t*) respectively. We consider time as a discrete variable, where each time step is a day. Similarly, at each time interval we track the proportion of humans within each compartment, where humans move from susceptible to infected according to a transmission rate, and humans move from infected to treated according to a treatment rate. Infected and treated humans return to the susceptible compartment according to recovery rates which not only vary according to whether treatment was received (as is standard for SITS models), but also on the proportion of the resistant pathogen within the host. This is where this model differs from other SITS models.

Unlike other SITS models, at each time interval we also track the pathogen population within humans, see Fig 1. Consequently, whereas compartmental models usually focus on the proportion of hosts within each compartment, here we focus on the overall Proportion of Resistant Pathogens (PRP), ***X***_*H*_ (*t*). Box 1 shows the time stepping process, where the population PRP is given by Eq. B, which depends on the proportion of infected *I*(*t*), Eq. B.1a, and treated humans *T* (*t*), Eq. B.1b; and the pathogen population within infected ***X***_*I*_ (*t*), Eq. B.2a, and treated humans ***X***_*T*_ (*t*), Eq. B.2b. The PRP is influenced by factors such as the probability of treatment, treatment effects, and most notably for this paper, the three competition dynamics:

- ↓ Transmission cost,
- ↓ Fitness cost in infected, untreated humans,
- ↑ Competitive release in infected, treated humans.

The arrows indicate the influence on increasing (↑) or decreasing (↓) resistance in the population. We will continue to use this notation throughout to remind the reader of the direction of the influence.

To demonstrate the effect of these competition dynamics within a single human, consider the example where a human is infected with a pathogen population where half the pathogen population is sensitive to treatment and the other half is resistant 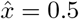 (*I_SR_* in Fig 1), and there are not multiple transmission events so an infected person cannot receive further infections. Initially, the proportion of the resistant pathogen may decrease due to a ↓ fitness cost associated with resistance. If treatment is received, the resistant pathogen proportion will increase as a result of clearance of sensitive pathogen, as well as growth of resistant pathogen due to additional resources available - this is ↑ competitive release. Finally, as this pathogen mix is transmitted onward, there may be a ↓ transmission cost which favours one class of pathogen over the other (the Between hosts dynamics shown in Fig 1). (From herein we assume that transmission only has a negative effect on resistance in the population, however this is not a restriction of the model.) There are more factors than the competition dynamics that affect 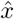, for example the more effective the drug, the more sensitive pathogen are cleared. Consequently, taking into account the many processes, a pathogen population transmitted to a human of 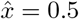 is unlikely to cause secondary infections with the same mix.

We discretise 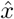 at *N* levels, and denote a particular level with *n* = 0, 1, … *N*. Throughout we take *N* = 10 (except in the schematic in Fig 1 where, for clarity of demonstration, *N* = 2). Therefore *x*_0_ corresponds to a pathogen population of only sensitive pathogen 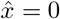, *x*_1_ corresponds to a pathogen population of 10% resistant pathogen 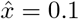, and so on, until *x_N_* = *x*_10_ which corresponds to a pathogen population of only resistant pathogen 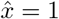. An example PRP over the course of an untreated infection, where the initial infection was *x_n_*, is presented in Fig 3. After the initial infection, the processes acting upon the PRP within hosts is continuous. The corresponding PRP over the course of a treated infection, where the infection on the day of treatment was *x_n_*, is presented in Fig 5.

The competition dynamics, and all other factors, are presented in the model schematics in Fig 1, which depicts the iterative process given in Box 1. The factors listed in the model schematics are considered in the global sensitivity analysis, where the variable of interest is the overall PRP within all humans - the population PRP. We note that some factors indirectly affect the population PRP via altering the proportion of humans within a compartment (such as treatment rate parameters), some factors directly impact the PRP but not directly impact the proportion of humans within a compartment (such as the competition dynamics), and some factors do both (such as drug efficacy).

Factors which affect the PRP within infected humans (not treated and treated) are generally captured by the distributions ***χ***_*I*_ (*a*) and ***χ***_*T*_ (*a*), where *a* ∈[1, *A_S_*] and *A_S_* is the maximum infection length. These are distributions of 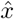, and are discretised at *N* + 1 levels where *χ_I,n_*(*a*) is the *n*th component of ***χ***_*I*_ (*a*), meaning it corresponds to 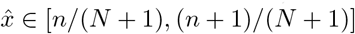. For example, *χ*_*I*,0_(*a*) is the component that corresponds to 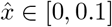 for *N* = 10. Similarly for *χ_T,n_*(*a*) as a component of ***χ***_*T*_ (*a*). At *a* = 1, ***χ***_*I*_ (*a*) and ***χ***_*T*_ (*a*) are uniformly distributed. If there were no within host dynamics, this distribution would remain uniform for all *a*, that is, there is no change to the PRP within hosts. However, as *a* → *A_S_* the distribution changes due to our model assumptions. Within untreated hosts, the distribution shifts so that the pathogen populations of mainly sensitive pathogen *χ*_*I*,0_(*A_S_*) dominate. Within treated hosts, the distribution shifts so that the pathogen populations of mainly resistant pathogen *χ_T,N_* (*A_S_*) dominate. Furthermore, the sum of the distribution on day *a* may be less than one, Σ_*n*_ ***χ***_*I*_ (*a*) ≤ 1 and Σ_*n*_ ***χ***_*T*_ (*a*) ≤ 1, since the infection length may be less than *A_S_* due to recovery. These differences from unity are referred to as the recovery rates, formally defined as,

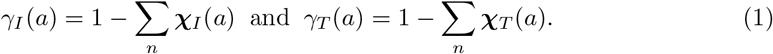

Note that the recovery explicitly depends on the *x_n_*, the discritised PRP within humans. The connection between our model assumptions which affects the PRP within hosts, and the distributions, are shown in Fig 3 and Fig 5 (the continuous PRP for different *x_n_*), and Fig 4 and Fig 6 (the distributions ***χ***_*I*_ (*a*) and ***χ***_*T*_ (*a*) discretised into *N* + 1 bins).

The rest of the Methods derive the model components in Box 1. After each process is introduced, we detail how each factor affects the population PRP, calculated by

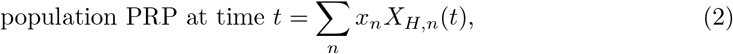

where *X_H,n_*(*t*) is the *n*th component of ***X***_*H*_ (*t*), as described above for *χ_I,n_*(*a*) being the *n*th component of ***χ***_*I*_ (*a*). Factors may indirectly affect the population PRP via altering the proportion of humans within a compartment, *I*(*t*) and *T* (*t*), or they may directly impact the PRP within humans, ***X***_*I*_ (*t*) and ***X***_*T*_ (*t*), or they may do both. Code is provided in Supporting Information.

#### Box 1: The SITS model presented in Fig 1 updates at each time interval according to the model equations described here. See Table 1 for further parameter descriptions and ranges.

The proportion of susceptible, infected and treated humans are denoted by *S*(*t*), *I*(*t*) and *T* (*t*) respectively. The overall Proportion of Resistant Pathogens (PRP) within infected and treated humans are denoted by ***X***_*I*_ (*t*) and ***X***_*T*_ (*t*) respectively. The overall PRP in all humans ***X***_*H*_ (*t*), which we refer to as the population PRP, is the weighted combination of the PRP in infected and treated humans,

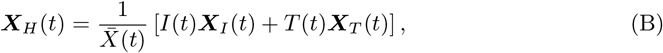

where 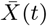 is a normalisation factor so that Σ_*n*_ ***X***_*H*_ (*t*) = 1. It is the population PRP from Eq. B that we track and examine in our results, which is updated at time *t*. First the proportion of infected and treated humans are updated, *I*(*t*) and *T* (*t*). Second the PRP within infected and treated humans are updated, ***X***_*I*_ (*t*) and ***X***_*T*_ (*t*). The details are:

**B.1a** The proportion of infected hosts *I* at time *t*, is the sum of all previous proportions of hosts infected, *βS*, that have not been treated (1 − *r*), and not recovered (1 − *γ_I_*),

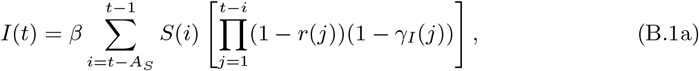

where *γ_I_* (*a*) is the recovery rate of infected hosts who have been infected for *a* days, and *A_S_* is the maximum infection length. (The infection length of an untreated infection which comprises sensitive pathogen only.)

**B.1b** The proportion of treated hosts *T* at time *t*, is the sum of all previous proportions of hosts infected, that were not treated for *k* days (nor recovered before treatment (1 − *γ_I_*)), but treated on day *k* + 1, and since this day, not recovered (1 − *γ_T_*),

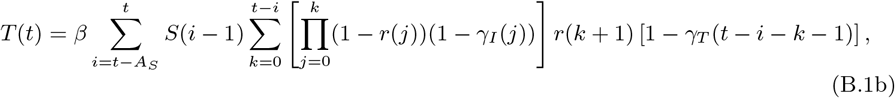

where *γ_T_* (*a* − *k* − 1) is the recovery rate of treated hosts who have been infected for *a* days, and treated on day *k* of their infection.

**B.2a** The distribution of resistant parasitaemia within infected hosts ***X***_*I*_ at time *t* depends on the distribution that hosts were infected with at the previous times ***X***_*V*_, and the dynamics that affect the distribution within hosts, encapsulated in ***χ***_*I*_,

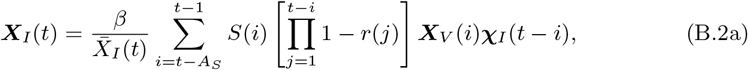

where 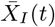 is a normalisation factor so that Σ_*n*_ ***X***_*I*_ (*t*) = 1.

**B.2b** The distribution of resistant parasitaemia within treated hosts ***X***_*T*_ at time *t* depends on the distribution that hosts were infected with at the previous times ***X***_*V*_, and the dynamics that affect the distribution within hosts, encapsulated in ***χ***_*I*_ and ***χ***_*T*_,

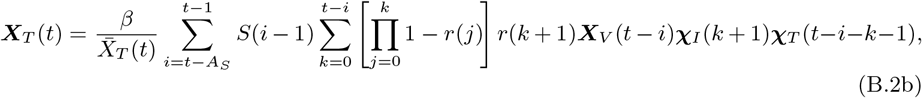

where 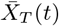 is a normalisation factor so that Σ_*n*_ ***X***_*T*_ (*t*) = 1. Distributions ***χ***_*T*_ and ***χ***_*T*_ capture the proportion which have not recovered 1 − *γ_I_* and 1 − *γ_T_*.

### The general transmission model

Humans are infected at a rate *β* per day, and we assume that humans infected on day *t* contract a PRP given by *X_V_* (*t*). The PRP that are transmitted from humans at time *t*, is given in Eq. B. If there are no transmission processes, then the PRP transmitted from humans is the same as the PRP transmitted to humans, *X_V_* (*t*) = *X_H_* (*t*), see Eq. B in Box 1. We allow the PRP to vary during the transmission process. Firstly, the transmission process may take *ℓ* days, such that ***X***_*V*_ (*t*) = ***X***_*H*_ (*t* − *ℓ*). For malaria, this represents the latent incubation period of mosquitoes being infected before becoming infectious, approximately 15 days, *ℓ* = 15. More importantly, we allow for a ↓ transmission cost of resistant infections, such that ***X***_*V*_ (*t*) = *f_H_* (***X***_*H*_ (*t* − *ℓ*)). The function *f_H_* allows us to account for infections where the PRP that is transmitted from humans may be different to the PRP that is transmitted to humans. For malaria, this allows us to incorporate the recombination process which occurs during the lag period. For details regarding modelling the ↓ transmission cost, see paragraph Competition dynamic: ↓ Transmission cost. The range for the transmission rate, *β* ∈ [0.0002, 0.0036], is chosen such that on a given day, the proportion of infected individuals ranges between ~ 2% and ~ 83%, when the other parameters are set to the default values in Table 1,

The model can be stochastic, so that on each day, humans are infected with a PRP drawn from the distribution *X_V_* (*t*). For large populations, this adds significant computation time, making long term analysis unfeasible. Thus, for our analysis to identify determinants of resistance dynamics, we have omitted stochasticity by assuming all humans infected on day *t* receive a PRP that corresponds exactly to *X_V_* (*t*).

As an addition, to account for multiple infection events, we shift the distribution within the infected populations (untreated and treated) by *β*. That is, the infection length is not increased (because of immune dynamics), but hosts experience a shift towards the distribution of the PRP that may be transmitted,

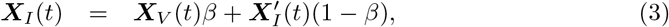

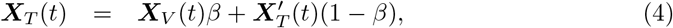

where 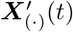 refers to the distribution within infected humans that is updated at each time interval according to Box 1. The prime notation is used only here for clarity of equations (3) and (4).

The transmission rate *β* directly affects the proportion of humans in the infected compartment, *I*(*t*) in Box 1. Therefore, it also indirectly affects the proportion of humans in the treated compartment, *T* (*t*) in Box 1. If allowing for multiple infection events, the additional shifts Eq. 3 and Eq. 4 means that the transmission rate *β* also affects the PRP within infected and treated hosts, ***X***_*I*_ (*t*) and ***X***_*I*_ (*t*) in Box 1.

### Between host transmissions

The PRP that are transmitted from humans at time *t*, ***X***_*H*_ (*t*), is given in Eq. B. As mentioned above, suppose there is a process where the infection that is transmitted from humans may be different to the infection that is transmitted to humans, ***X***_*V*_ (*t*) = *f_H_* (***X***_*H*_ (*t* − *ℓ*)). This allows us to model potential costs to the onward transmission of resistant infections.

#### Competition dynamic: ↓ Transmission cost

At a given time the human population has a certain distribution of PRP, comprising some infections which are all sensitive, some are all resistant, and some are mixed infections. Then, due to a transmission cost, the distribution that is transmitted to humans, after a period *ℓ* ≥ 0, is altered by the following,

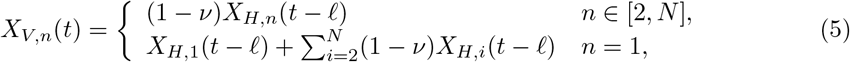

where *ν* ∈ [0, 1], *X_V,n_*(*t*) is the component of ***X***_*V*_ (*t*) corresponding to PRP *x_n_*, and *X_H,n_*(*t*) is the component of ***X***_*H*_ (*t*) corresponding to PRP *x_n_*. By varying *ν* we can alter the cost dynamics such that when *ν* = 0 there is no cost to transmission, and when *ν* = 1, a resistant infection is impossible to transmit, representing an infinite transmission cost to resistance.

For malaria, recombination of sexual parasites within the mosquito is a transmission cost. The malaria parasite reproduces sexually within a mosquito, but not humans. Often several mutations are required for resistance, which may be split during sexual reproduction. Consequently, the next generation of infection is no longer resistant. For example, consider a mosquito carrying one sensitive infection, and one resistant where resistance occurred due to two mutations on the chromosome. After recombination, the two mutations may be split so that the mosquito transmits two infections which each only have one mutation (two partially resistant infections). In the case of one mutation only (such as resistance to chloroquine [43, 44]) recombination has no effect, *ν* = 0 in Eq. 5. Whereas, in the case of infinite mutations, *ν* = 1. However, chloroquine resistance (single locus) and resistance to combinations therapies, such as sulfadoxine-pyrimethamine and artemisinin-based therapies, (multi-locus) exhibit similar spread at the population level. For this reason, [45], who present a nested model (within host and between host) for the spread of drug-resistant malaria, exclude recombination. We include recombination here to determine whether it is more important in some settings than others.

If transmission does not come at a cost to resistant pathogens, but as a gain, then transmission competition dynamics favour resistant pathogens. This gives the following alternative to Eq. 5,

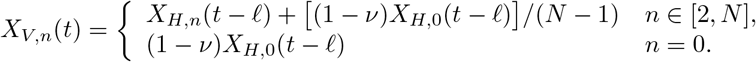

For malaria this is less likely, and thus we did not consider this scenario.

The recombination factor *ν* does not directly affect the proportion of hosts who are infected, nor treated. It affects the PRP within newly infected hosts, ***X***_*I*_ (*t*) in Box 1, via ***X***_*V*_ (*t*). Therefore, it also indirectly affects the proportion of humans in the treated compartment, ***X***_*T*_ (*t*) in Box 1.

### Within infected but untreated humans

With a new infection each human is transmitted a pathogen population where *x_n_* (*n* = 1, 2,… *N*) proportion is resistant. The number of days since a host was initially infected is *a* ≥ 0. A host is expected to be infected for *A* days, which we assume depends on *x_n_* such that,

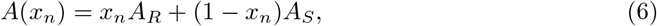

where *A_R_* and *A_S_* are the expected infection length, in days, of resistant and sensitive infections in an untreated human. Eq. 6 allows the total average length of infection to vary if resistant infections incur a fitness cost, meaning that resistant pathogens are weaker and potentially shorter (see Competition dynamic: Fitness cost paragraph below). Therefore, within untreated humans, resistant infections have a shorter infection length, *A_R_ ≤ A_S_*. Note that the maximum infection length, *A_S_*, is only incurred by untreated humans carrying only sensitive pathogen.

#### Competition dynamic: ↓ Fitness cost

The ↓ fitness cost modelled here not only accounts for reductions in the infection length of resistant infections, but in the case of mixed infections, the PRP is suppressed as the sensitive pathogen dominates. Therefore, a human infected with both pathogens may become a host to only sensitive pathogens (if untreated). We assumed the PRP during the infection period *a* ∈ [0, *A_S_*] follows a function,

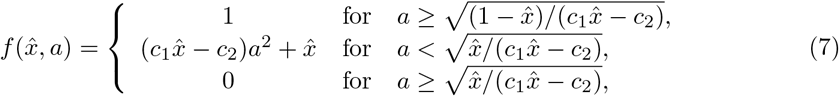

where *c*_1_, *c*_2_ ≥ 0 are constants indicating the advantage that sensitive have over resistant infections. When *c*_1_ = *c*_2_ > 0 sensitive infections dominate (this is the most realistic scenario [38, 46]), when *c*_1_ > 0, *c*_2_ = 0 resistant infections dominate, and when *c*_1_ = *c*_2_ = 0, there is no advantage in either direction.

In our analysis, we consider the cases where there are no fitness costs, and where sensitive infections dominate, *c* = *c*_1_ = *c*_2_ ≥ 0. We assume the reduction of the infection length of resistant infections *A_R_* is related to the length of sensitive infections *A_S_* by

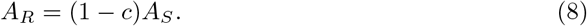

The infection lengths *A_S_* and *A_R_* affect the recovery rate *γ_I_* (*a*) (determined from the distribution ***χ***_*I*_ (*a*), see Eq. 1 and Fig 4). Consequently *A_S_* and *A_R_* directly affect the proportion of humans in the infected compartment, *I*(*t*) in Box 1. Therefore, they also indirectly affect the proportion of humans in the treated compartment, *T* (*t*) in Box 1. Additionally *A_S_* and *A_R_* directly affects the PRP within untreated hosts and treated hosts, ***X***_*I*_ (*t*) and ***X***_*T*_ (*t*) in Box 1, again via ***χ***_*I*_ (*a*). Note that the dynamics, Eq. 7 and Eq. 8, affect mixed infections only.

To summarise the factors which affect the PRP in infected, untreated, hosts, ***X***_*I*_ (*t*): Overall, at each time there are *I*(*t*) infected hosts who were infected on different days (so with different initial *x_n_*), and have a new *x_n_* at time *t* due to the fitness cost *c*.

Therefore, the PRP in infected hosts, ***X***_*I*_ (*t*), depends on

- the PRP transmitted ***X***_*V*_ (*t*) for each day previously, up to *t* − *A* days prior.
- the continued change to the PRP due to the within host fitness cost, Eq. 7 and Eq. 8, which are captured by ***χ***_*I*_ (*a*).

### Treatment

Unlike standard SITS models, the treatment rate *r*(*a*) is not constant, but instead dependent on the maximum probability of being treated *r_x_*, and the period in which treatment is possible 1*/α*, see Fig 2. This is to capture that treating early in the infection treats a different pathogen mix compared to later in the infection. Moreover, it allowed us to incorporate non symptomatic infections by assuming that if an infection was untreated for a given period, then treatment would not occur at all.

**Figure 2.**
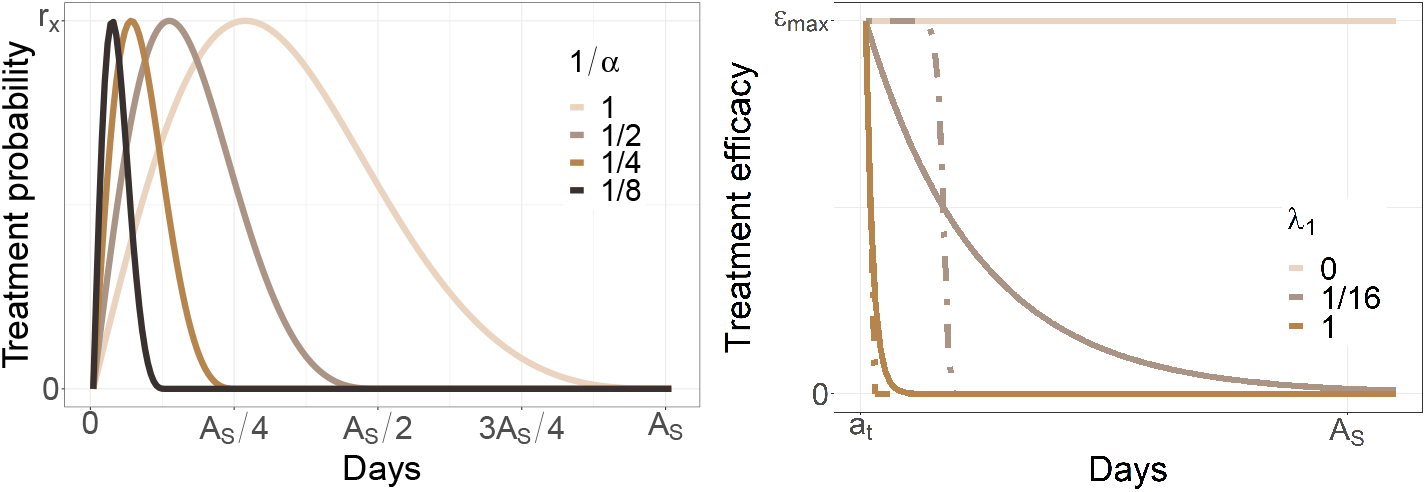
Illustration of treatment probabilities and treatment efficacy. Left: The probability of treatment for different values of *α*, as a function of the infection time, is modelled using a skewed sine curve, see Eq. 9. The maximum probability is *r*(*x*). The day that this maximum occurs, and the last possible day of treatment, are defined by *α*, which varies here between 1 and 8. Right: The drug concentration within a treated host, as a function of the infection time, is modelled using a Weibull decay like function, see Eq. 10. The concentration begins at *ϵ_max_* and decays according to two shape parameters: the decay rate *λ*_1_ and a delay before decay *λ*_2_. When *λ*_2_ = 1 there is no delay and the decay is exponential at rate *λ*_1_ (solid line). The maximum delay we allow is *λ*_2_ = 19 (dot-dashed line).

The treatment rate over the maximum possible infection length *A_S_* is captured using a skewed sine curve,

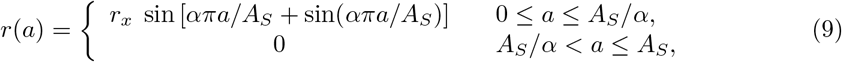

where *r_x_* is the maximum probability of being treated, and *α* ∈ [0, *A_S_*] reflects the last possible day of treatment. The minimum value, *α* = 0, indicates no treatment, and the maximum value, *α* = *A_S_*, indicates that treatment is possible over the whole infection length.

To parameterise *r_x_* we consider that the probability of receiving treatment is between 5% and 80% in a two week period, with an average at 40% [47]. Therefore, for a given human, the probability of treatment over a two week period can be represented as,

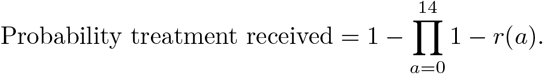

This guides reasonable values of *r_x_* such that if the probability of treatment in two weeks is 5%, 40%, or 80%, then *r_x_* is 0.01, 0.08, and 0.24 respectively.

The treatment rate parameters *r_x_* and 1*/α* directly affects the proportion of humans moving from the infected compartment to the treated compartment (see Box 1). Specifically, on day *t* the proportion of infected hosts *I*(*t*) depends on the probability of not being treated previously, that is ∏_*j*_ (1 − *r*(*j*)), and the proportion of treated hosts *T* (*t*) depends on the probability of not being treated for *k* days previously, and then treated on day (*k* + 1), that is 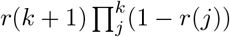. The *r*(*j*) are determined by Eq. 9.

### Drug concentration

The maximum concentration of the drug in the host *ϵ_max_* is assumed to be on the day of treatment *a_T_*. After this day we allow the concentration to decay at a rate *λ*_1_, which reflects the different half lives of drugs. Initially the decay is inhibited by *λ*_2_ which reflects a delay before a decay of the drug concentration within the host. We model the potential decay of drug efficacy *ϵ*(*a*) as a Weibull decay function, given by

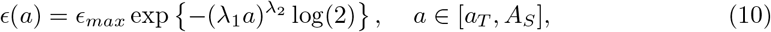

see Fig 2, and the green shaded region in Fig 5. When *λ*_2_ = 1, the decay rate is exponential, with decay *λ*_1_, representing a single use drug that decays immediately. As *λ*_2_ increases, the decay process determined by *λ*_1_ is delayed by several days. When *λ*_1_ = 0, the drug concentration does not decay, and when *λ*_1_ = 1, the complete decay is nearly immediate.

The drug concentration parameters, *ϵ_max_*, *λ*_1_ and *λ*_2_, do not directly affect the proportion of hosts who are infected, however they do directly affect the proportion of hosts who are treated, *T* (*t*) in Box 1. That is, the proportion of humans leaving the treated compartment is dependent on the time a treated host takes to recover, *γ_T_* (*a*), (see Recovery of treated humans paragraph below). Additionally, the drug concentration parameters directly affects the PRP within treated humans, ***X***_*T*_ (*t*) in Box 1. This is because Eq. 10 affects the PRP within treated hosts, see Fig 5, which affects the distribution ***χ***_*T*_ (*a*), see Fig 6.

### Within treated hosts

When a human is treated on day *a_T_* of their infection, with a drug that is *ϵ_max_* effective, then *ϵ_max_* of the sensitive pathogen are removed on day *a_T_*, and thus increases the PRP to 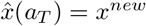. The new PRP is

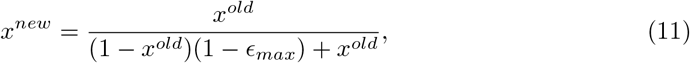

where *x^old^* is the proportion just before treatment, and *ϵ_max_* is the efficacy of the drug. To demonstrate the intuition of Eq. 11, consider the following example: at the time of treatment 60% of the pathogen population within an infected host is resistant, *x^old^* = 0.6. This person is treated with a drug that is 75% effective, *ϵ_max_* = 0.75, so *ϵ_max_*(1 − *x^old^*) = 30% of the pathogen are removed,

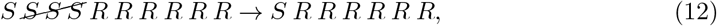

and thus *x^new^* = 0.86. The development from *x^old^* to *x^new^* is independent of competition dynamics. Notably, even without ↑ competitive release, treating humans removes sensitive pathogen from the population. Consequently, over time, only resistant pathogens persist.

Within a treated host, after this initial clearance of *ϵ_max_* of the sensitive pathogen (Eq. 11), the PRP depends on the recovery rate *γ_T_* (*a*), the ↓ fitness cost *c* which continues to suppress the PRP, the concentration of drug which remains in the body *ϵ*(*a*), and ↑ competitive release. The affect that *ϵ_max_* has on the time iterative process given in Box 1 was described at the end of the Drug concentration paragraph above.

#### Competition dynamic: ↑ Competitive release

Whilst the host remains infected, ↑ competitive release means that with the clearance of sensitive pathogen, resistant pathogens have more resources available - they are released from competition. We model this process by first considering that at a maximum, the proportion of sensitive pathogen that is cleared by treatment (in the example in Eq. 12, this is 30%), are completely replaced by the resistant pathogen. Therefore, the PRP within a treated host evolves such that

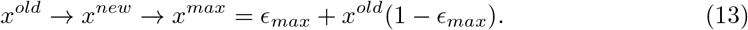

Continuing the example in Eq. 12,

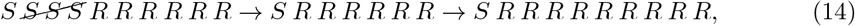

which corresponds to *x^old^* = 0.6, *x^new^* = 0.86 and *x^max^* = 0.9. In our model we assume that competitive release occurs incrementally, each day, by *κ* such that *κ* = 0 means *x^max^* is never reached as there is no competitive release, and *κ* = 1 means that competitive release is so strong that *x^new^* = *x^max^*. We consider *κ* ∈ [0, 0.2], since as we later observe, there is no further insight to be gained by increasing this range.

To demonstrate the effect of treatment on the PRP within hosts, suppose that at the time of treatment *x^old^* = {0, 0.1,…, 1}. Upon treatment, the PRP jumps up according to Eq. 11, see the PRP at time *a_T_* in Fig 3. Then the sensitive pathogen is suppressed according to *ϵ*(*a*) (the shaded green), keeping the PRP elevated. Once the drug has cleared, the ↓ fitness cost can again dominate, driving the PRP down. This reduction is hindered according to the ↑ competitive release, which varies for Fig 3(A)–Fig 3(C).

**Figure 3.**
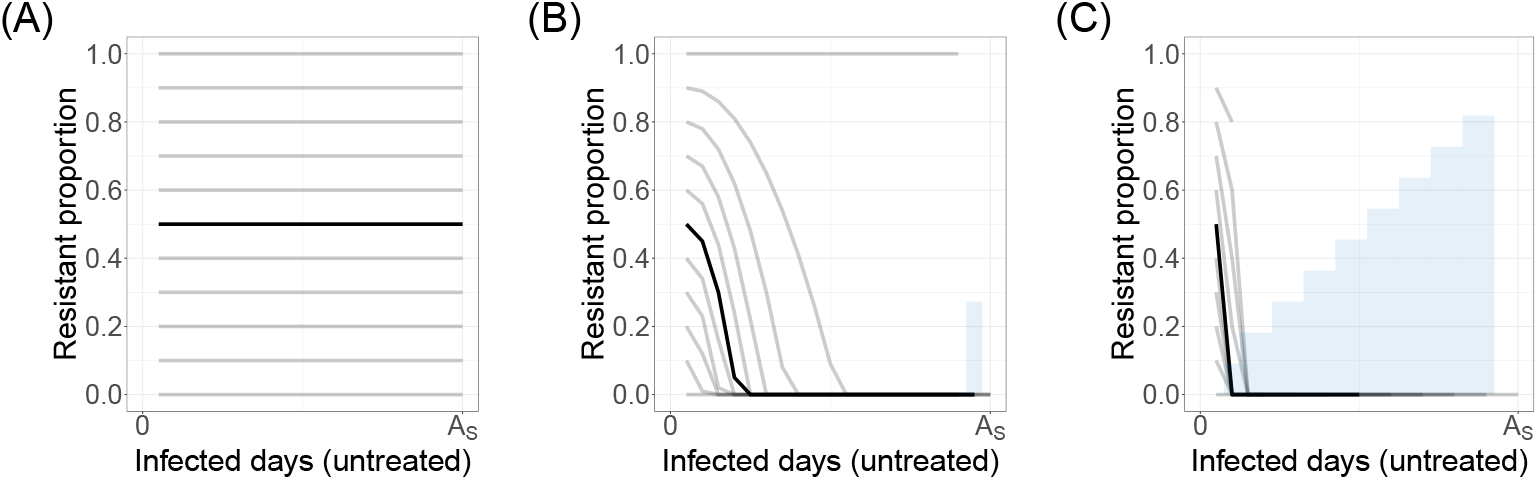
Example PRPs, within untreated humans, varying over the infection length. The initial PRP is *x_n_* = 0, 0.1,…, 1 (grey lines). A patient infected with 50% PRP (*x*_5_ = 0.5) is highlighted in black. PRP changes over the infection length according to the fitness cost (Eq. 7). The infection length depends on the initial PRP (Eq. 6) and the fitness cost (Eq. 8). (A) No fitness cost *c* = 0. (B) Some fitness cost *c* = 0.1. (C) Maximum fitness cost *c* = 1. The proportion which recover before the maximum infection length *A_S_* are shown in blue.

**Figure 4.**
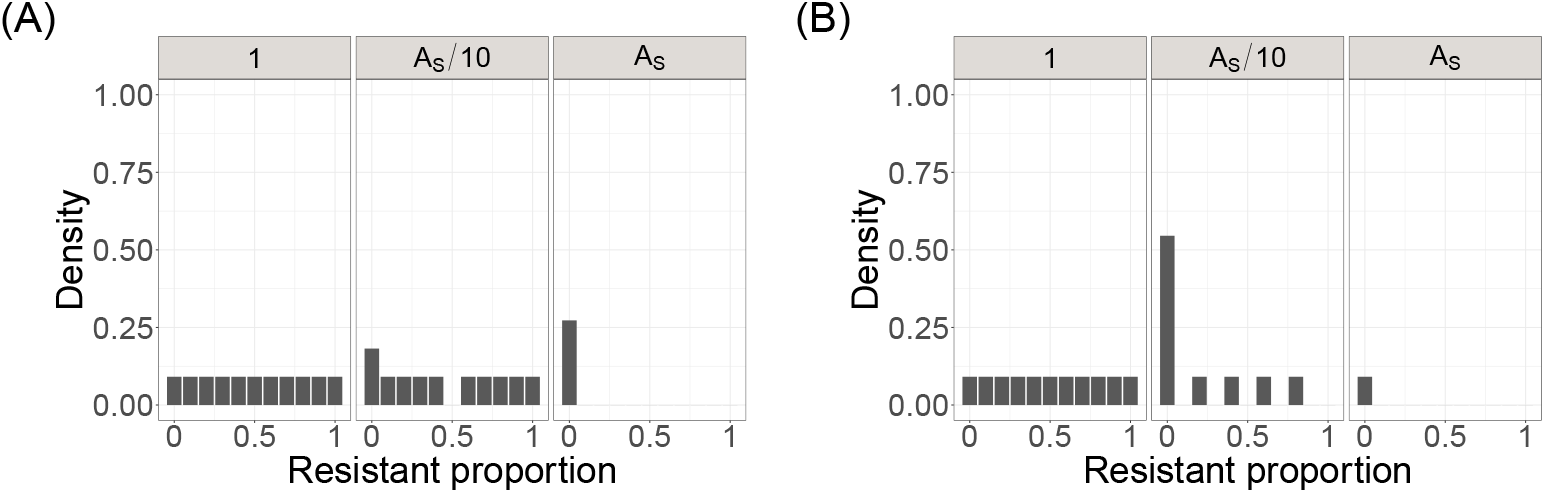
The distributions *χ*_*I*_ (*a*) from the PRP. The panels correspond to the PRP shown in Fig 3B and Fig 3C at the time of infection, *a* = 1, some time after infection, *a* = *A_S_/*10, and at the maximum infection length *a* = *A_S_*. (A) Some fitness cost, *c* = 0.1. (B) Maximum fitness cost *c* = 1. The case for no fitness cost (Fig 3A) would remain as a uniform distribution for all *a*. The density is less than one as *a* → *A_S_* to represent the recovered proportion (shown in blue in Fig 3B and Fig 3C).

In our model we assume that ↑ competitive release *κ* does not affect the recovery rate. Therefore ↑ competitive release does not affect the proportion of humans moving from the treated compartment. However, it affects the PRP within treated humans, ***X***_*T*_ (*t*) in Box 1, via ***χ***_*T*_ (*a*) see Fig 6.

### Recovery of treated humans

The time taken to clear the infection varies proportional to the treatment efficacy *ϵ_max_*. When *ϵ_max_* is close to one, sensitive pathogen are cleared within a few days, and when *ϵ_max_* = 0, sensitive pathogen remain unaffected by treatment. More generally, accounting for a mixed pathogen population, treatment administered to a human on day *a_T_* ∈ [0, *A_R_*] will continue to be infected for

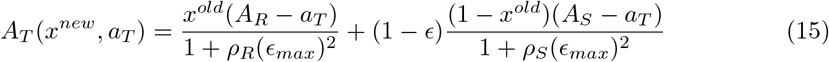

days, where *ρ_R_* ∈ [0, 1] and *ρ_S_* ∈ [1, 19] are scaling variables. When *ρ_R_* = 0, resistant infections remain unaffected by treatment, so that the infection length of a fully resistant infection would remain unchanged; and when *ρ_R_* = 1, the infection length of a fully resistant infection is halved, which we assume is the upper bound in reduction of infection length for a fully resistant infection. This same bound is assumed to be the lower bound for the reduction of infection length for sensitive infections. That is, when *ρ_S_* = 1 the infection length of a sensitive infections is halved. As an upper bound, we assume that the infection length of fully sensitive infections, can at most, be divided by 20. These limits are chosen so that when drawn randomly, treated sensitive pathogen will always clear quicker than treated resistant pathogen.

When the treatment is ineffective, *ϵ_max_* = 0, Eq. 15 reduces to Eq. 6, the infection length for an untreated infection. Therefore this is no discontinuity in our assumption of the effect of treatment on infection length. Also note that if the infection is all sensitive, and the treatment is fully effective, *ϵ_max_* = 1, then the infection is immediately cured.

We assumed that the recovery rate of treated humans is updated from the recovery rate of infected humans due to the maximum drug efficacy *ϵ_max_*, and the PRP at the time of treatment *x^old^* (not competition dynamics). Consequently, although treatment initially shifts the within host PRP so that there is a larger PRP, *x^new^* ≥ *x^old^*, humans carrying a high proportion of sensitive pathogen, at the time of treatment, recover quickly. That is, as *a_T_* → *A_S_*, treated humans which have not recovered have a high PRP, see Fig 5.

**Figure 5.**
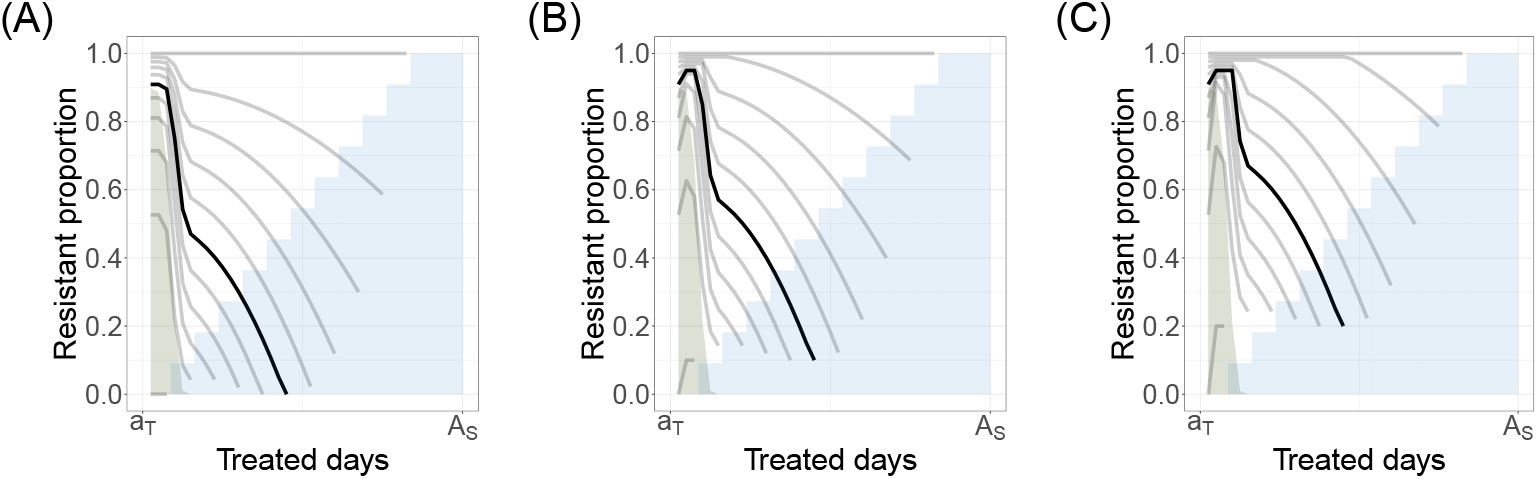
Example PRPs, within treated humans, varying over the remaining infection length. The PRP at the time of treatment is *x_n_* = 0, 0.1,…, 1 (grey lines). A patient with 50% PRP (*x* = 0.5) at the time of treatment is highlighted in black. PRP changes over the remaining infection length according to the fitness cost (Eq. 7), the treatment efficacy (Eq. 11), and competitive release (Eq. 13). The remaining infection length depends on the reduction due to treatment (Eq. 15). Recovery before the maximum infection length *A_S_* is shown in blue. The drug efficacy (Weibull decay) is shown in green. (A) No competitive release, *κ* = 0. (B) Some competitive release *κ* = 0.1. (C) Maximum competitive release, *κ* = 0.2.

**Figure 6.**
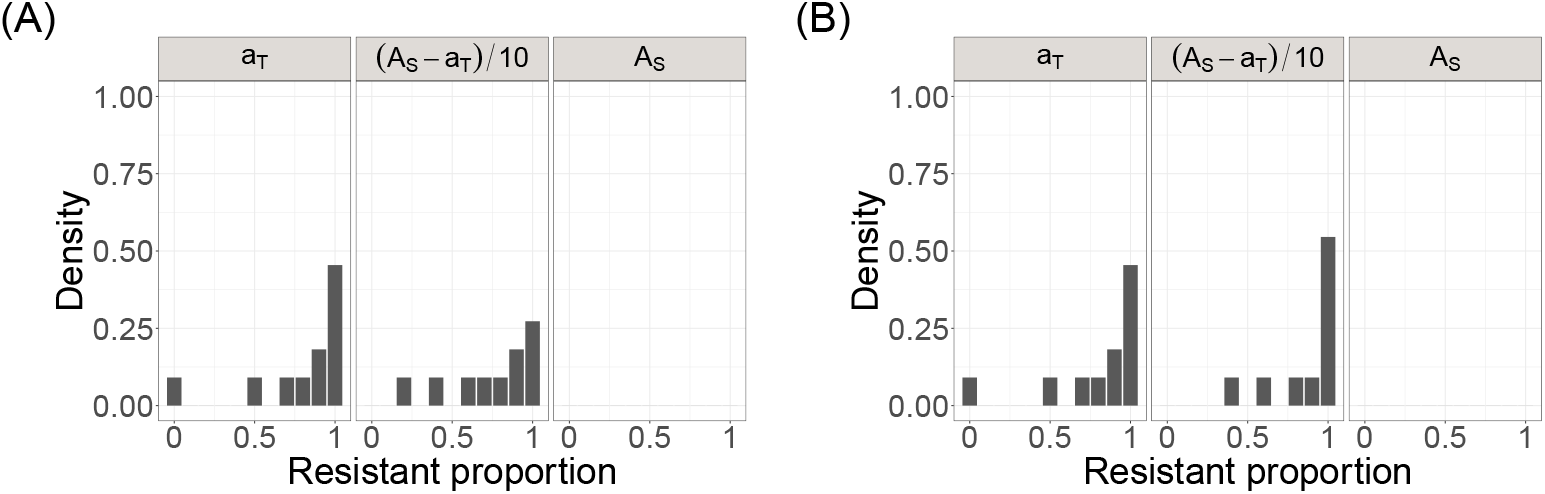
The distribution *χ*_*T*_ (*a*) from the PRP. The panels correspond to the PRP shown in Fig 5A and Fig 5C at the time of infection, *a_T_* = 1, some time after treatment, *a* = (*A_S_* − *a_T_*)/10, and at the maximum infection length *a* = *A_S_*. (A) No competitive release, *κ* = 0. (B) Maximum competitive release, *κ* = 0.2. The case for some competitive release (Fig 5B), *κ* = 0.1, is between these two examples. The density is less than one as *a* → *A_S_* to represent the recovered proportion (shown in blue in Fig 5A and Fig 5C).

The parameters that reduce the length of resistant *ρ_R_* and sensitive *ρ_S_* pathogen induced infections directly affect the proportion of humans who are treated, *T* (*t*) in Box 1, via the recovery rate *γ_T_* (*a*). Additionally, whilst the reduction in infection lengths do not directly shift the PRP within treated humans (in the same way that drug efficacy and competition dynamics do), by clearing sensitive pathogens quicker, *ρ_R_* and *ρ_S_* affect ***χ***_*T*_ (*a*) and thus ***X***_*T*_ (*t*) in Box 1.

### Addition of resistance

In our model 0.1% resistance is added to the population after the system has settled to a steady state. This could represent de novo mutations, or new resistant infections, entering the population. Following this event the proportion of resistance may increase due to treatment clearing sensitive pathogens, and due to ↑ competitive release. That is, further de novo mutations are not included because (i) the mutation rate is typically very small, and (ii) repeatedly shifting the proportion of resistant pathogen in manner that is not connected to other aspects of model provides little insight into the tug of war of the competition dynamics. Nonetheless, just as we add 0.1% resistance once, it is straightforward to add any given percentage of resistance, at any rate, throughout the simulation, and thus incorporate a mutation rate. Alternatively, the effect of *κ* may be interpreted as the effect of both ↓ competitive release and de novo mutation combined. This alternative interpretation, may be suited for cases like *M. tuberculosis*, where de novo evolution is frequent.

### Sensitivity analysis

The model is run for 3000 different scenarios, where each scenario is defined by the unique set of parameter values drawn randomly from the ranges listed in Table 1. After three years, resistance is added, and the simulations are stopped at 20 years. The population PRP at 20 years is recorded for each simulation, and we refer to this as a measure of the spread of resistance.

A simulation for 20 years (7300 days), with *N* = 10, takes approximately 16 minutes on a standard PC. The memory required for a single simulation is related to the days in the simulation, and the discretisation, by order *O*(totalDays^2^ × *N*).

#### Establishment

We split the scenarios into two based on the population PRP at 20 years: those where the population PRP is above zero (establishment), and those where the population PRP is zero (no establishment). To identify features that distinguish between these two scenario classes, we compare the distributions of the parameter values. A straightforward comparison is the difference in the (normalised) median of the parameter values under each class. A positive difference indicates that a large parameter value is associated with establishment, and a negative difference indicates that a large parameter value is associated with non-establishment. Plotting the two distributions provides insight that goes beyond a linear comparison. Note that the PRP threshold of zero for establishment can easily be changed.

#### Spread

Given establishment, we refer to the spread of resistance as the population PRP at 20 years. Taking only the scenarios where resistance occurred, we conduct a sensitivity analysis using the tgp R package [49, 50]. The package approximated the relationship between the 12 inputs and output (PRP) using a Gaussian Process with jumps to a limiting linear model (bgpllm). The model was calibrated to the scenarios where resistance established. To verify the fit of the calibration, we made predictions for additional scenarios, which were not included in the sensitivity analysis.

## Results

Our novel model described in Box 1, with details in the Methods section, tracks the PRP at the population level. The population PRP is influenced by factors such as the probability of treatment, treatment effects, and most notably for this paper, the three competition dynamics:

- ↓ Transmission cost,
- ↓ Fitness cost in infected, untreated humans,
- ↑ Competitive release in infected, treated humans.

The arrows indicate the influence on increasing (↑) or decreasing (↓) resistance in the population. The model is an SITS model, with several time dependent processes incorporated, resulting in 12 different input parameters, see Fig 1. These 12 parameters vary within the bounds in Table 1 so that each of the 3000 simulations are different.

To demonstrate the effects of the competition dynamics, an illustrative example is shown. We then present the results from the sensitivity analysis to determine which factors drive resistance establishing, followed by the results from the sensitivity analysis to determine which factors drive the spread of resistance, given that it has established. Separating establishment and spread highlights that they are not necessarily driven by the same factors.

### Illustrative example

A single simulation of the model was run for three years before resistance was added, as shown in Fig 7. In this example, without any competition dynamics, resistance increases since sensitive pathogens are continually being cleared, Fig 7(A). When ↑ competitive release is included, resistance dominates the population quicker, Fig 7(B). However, when all the dynamics are included, resistance can be suppressed, Fig 7(C).

**Figure 7.**
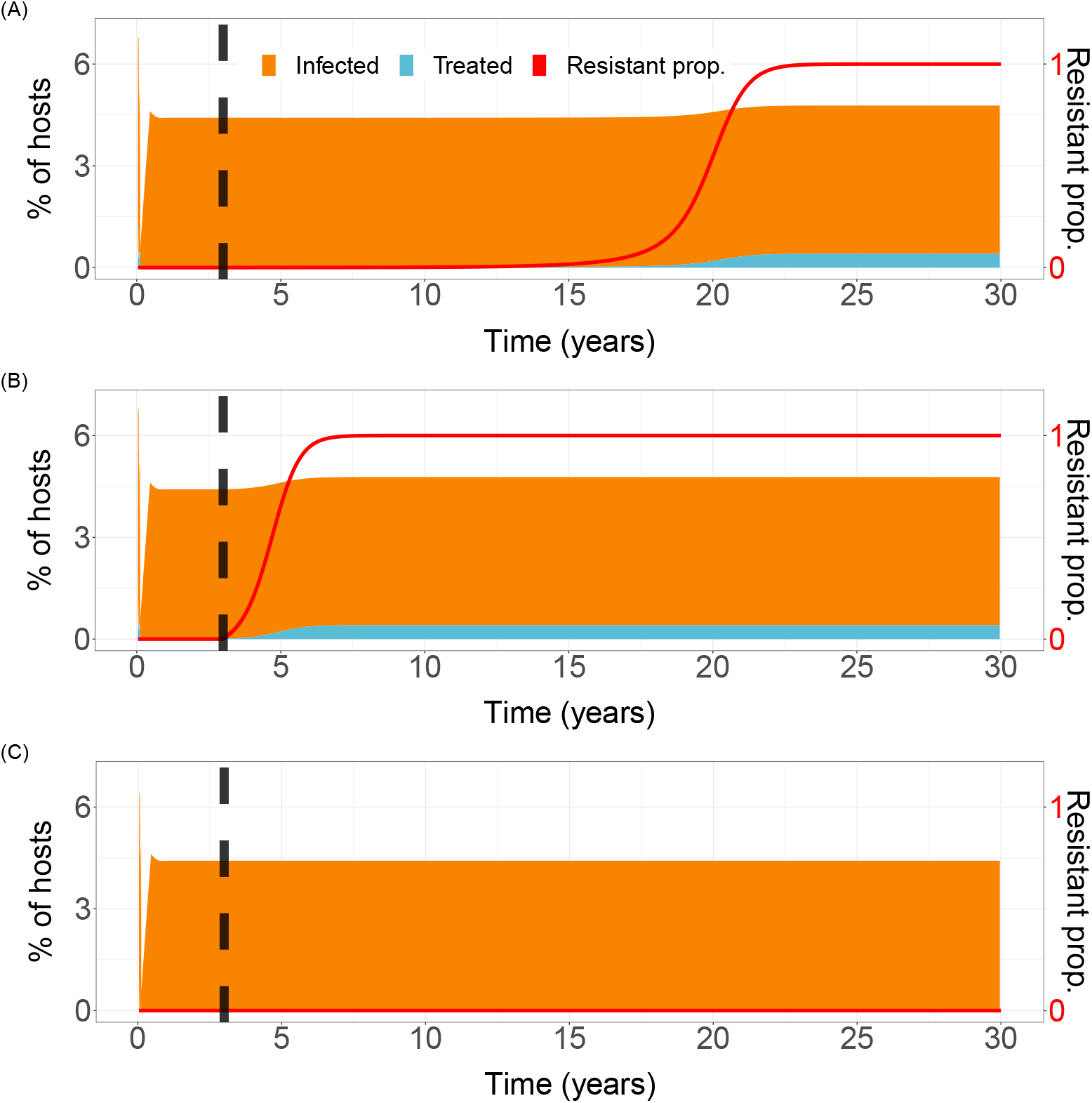
Demonstration of the proportion of infected and treated hosts, and the population PRP, varying over the 20 year simulation period. Parameters as in Table 1 and competition dynamics as follows. (A) No competition dynamics, *κ* = *c* = *ν* = 0. (B) ↑ Competitive release only, *κ* = 0.2. (C) ↑ Competitive release, ↓ fitness cost, and ↓ transmission cost, *κ* = 0.2, *c* = 1, *ν* = 1. The black dashed line, at three years, is when resistance is included.

### Establishment

Establishment occurred in 44% of our scenarios (1321 of 3000 scenarios). To obtain a detailed understanding of how parameters influence resistance establishing, we compare the distributions of the model parameters which led to resistance establishing with the distributions of the model parameters which did not lead to resistance establishing. Precisely, for a given parameter, we used the Komogorov-Smirnov test to identify when the distributions of the model parameters is significantly different when resistance establishes and when it does not (one-tailed 5% significance level where both directions were tested independently). This led us to conclude that the following six parameters have a strong association with establishment (in order of significance): ↑ competitive release, the maximum treatment efficacy, the proportion of untreated infection length in which treatment is possible, the maximum treatment probability, ↓ fitness cost, and reduction of infection length of sensitive infections when treated. In these cases we visually compare the two distributions, see Fig. 8, and discuss the differences. The p-values are in the caption of Fig. 8.

**Figure 8.**
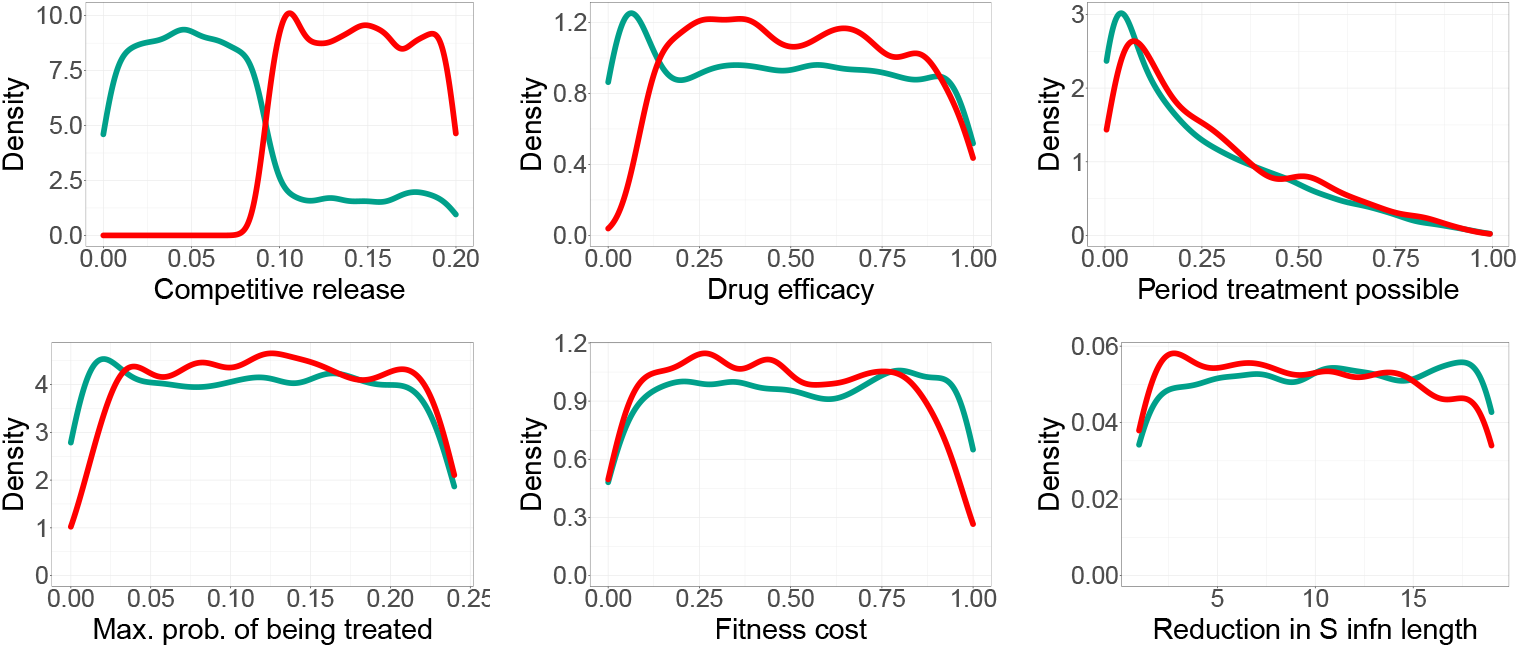
The distribution of parameters for the 1321 simulations where resistance established (red), and the 1679 simulations where it did not (green). The parameter is indicated by the *x*-axis label. These six parameters proved to have distributions that were significantly different using Komogorov-Smirnov test. The corresponding p-values are 0, *O*(10^−34^), *O*(10^−15^), *O*(10^−8^), *O*(10^−7^), 0.0026

By far, ↑ competitive release has the strongest association with establishment. Comparing the distributions (Fig. 8) we observe that ↑ competitive release almost has a switch-like function: when it is off (*κ* < 0.0925), establishment does not occur, and when it is on (*κ* > 0.0925), establishment occurs under 81.3% of the scenarios. Therefore, if the resistant proportion within hosts do not have the possibility to increase by 9.25% each day that the drug concentration is present, resistance cannot establish. The fact that resistance does establish in real life suggests that ↑ competitive release does occur.

Following ↑ competitive release, the next three significant parameters are treatment parameters: the maximum treatment efficacy, the proportion of untreated infection length in which treatment is possible, and the maximum treatment probability. The first from this list, the maximum treatment efficacy, describes the proportion of sensitive pathogen cleared within hosts, and ranges from zero to perfect. Therefore, when treatment efficacy is very low, it is similar to no treatment at all, and thus resistance is unlikely to establish. Comparing the distributions (Fig. 8), we observe that when the efficacy is very high, say above 90% clearance, it is slightly harder for resistance to establish. The other two treatment parameters are unique to this model because of our function to allow for a non-constant treatment rate (see Fig. 2 for further explanation). To capture the fact that treatment is more likely at the beginning of the infection, our function for the treatment rate is summarised by two parameters: the period treatment is possible and the maximum probability of being treated. This representation of a non-constant treatment rate, and our comparison of the distributions, highlights that when treatment is provided early in the infection, resistance is less likely to establish. In fact, we find this timing to be a stronger driver of the establishment of resistance than the maximum treatment probability.

When the ↓ fitness cost is high, resistance is significantly less likely to establish (Fig. 8). However, the ↓ fitness cost needs to be greater than approximately 0.8, which is an extreme ↓ fitness cost. Recall that a fitness cost of 1 implies that the proportion of untreated resistant pathogens are completely removed / replaced almost immediately by sensitive pathogens, see Fig. 3. We note that the other cost of resistance, the ↓ transmission cost, was not significantly different under both scenarios (p-value 0.377).

Our results show that when the reduction in sensitive infection length is greater than a given value (*ρ_S_* > 10), resistance is less likely to establish. That is, resistance is less likely to establish when sensitive infections are more responsive to treatment.

Quantifying the influence of parameters on establishment by comparing their distributions does not separate the single and total effects of parameters. We expect interactions to play a significant role because we designed the model to include known dependencies between parameters. For example, the effect of the ↓ transmission cost is dependent on the transmission rate. The transmission rate, the treatment rate, and recovery of non-treated humans, interact with all other factors (Box 1). That is, they impact the proportion of humans within infected and treated compartments, which indirectly, but fundamentally, affects the population PRP. The single and total effects of the parameters are considered separately in the next subsection. We performed a sensitivity analysis on the scenarios where resistance was established in the population.

### Spread of resistance given establishment

In the 1321 scenarios where establishment occurs, ↑ competitive release has a negligible effect on the population PRP, Fig 9. The population PRP is only sensitive to the drug efficacy, the maximum probability of being treated, the reduction in the length of sensitive infections due to treatment, and the period treatment is possible - in this order when considering first order effects. The total effects shows that the sensitivity of all variables increases, indicating a high level of interaction. Consider the reduction in the sensitive infection length upon treatment, and the period that treatment is possible: the sensitivity of these two parameters depends on whether one considers the single or total effects. Therefore, due to interactions, the period that treatment is possible has a greater total effect.

**Figure 9.**
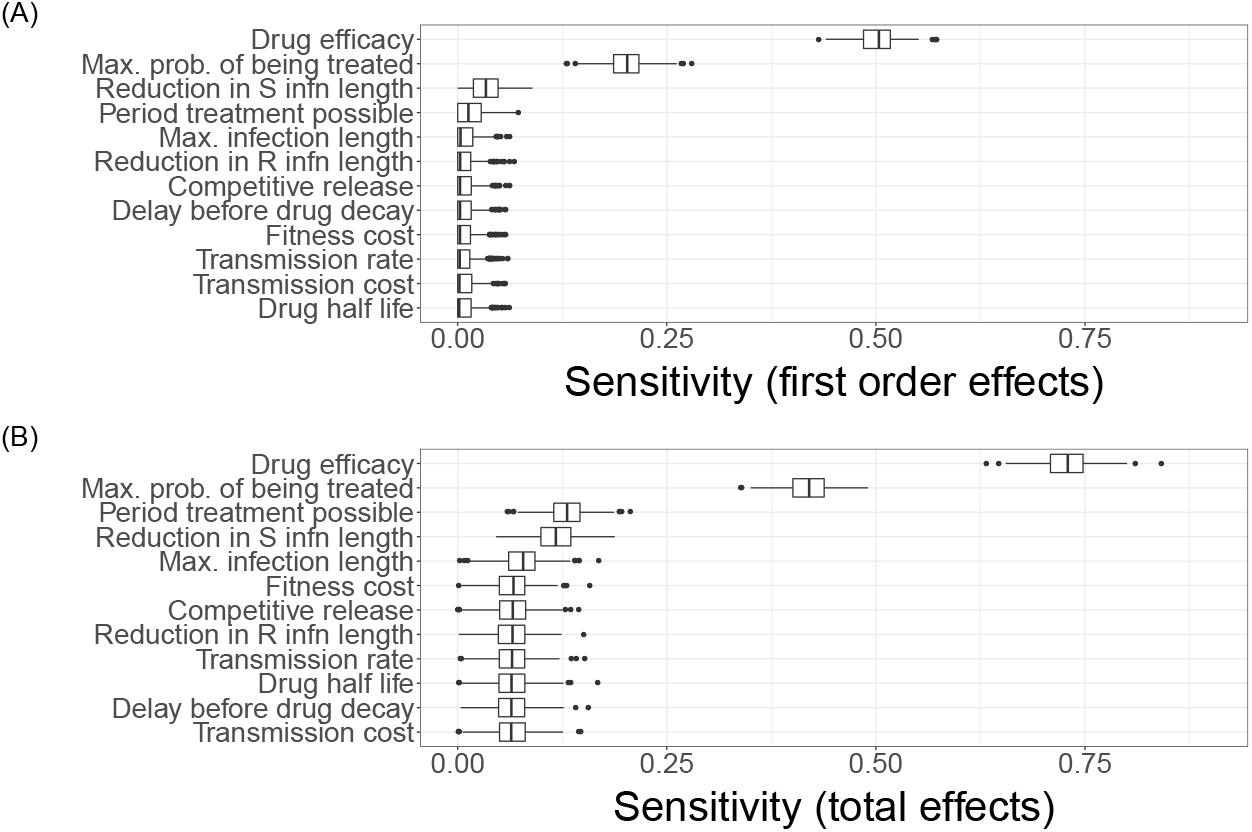
The sensitivity indices of parameters on the population PRP at 20 years, from 1321 simulations where resistance established. (A) First order effects (medians sum to 0.77). (B) Total effects (medians sum to 1.93).

Previously, we found that the period which treatment is possible was more strongly associated with establishment than the maximum probability of receiving treatment. Now, when examining only the scenarios where establishment occurs, this order is inverted such that the population PRP is more strongly influenced by the maximum probability of receiving treatment than the period that treatment is possible. Fig 10 shows that as the maximum probability increases, so does the population PRP, whereas the relationship with the period that treatment is possible is less straightforward. Generally, when treatment is possible over a larger portion of the infection period, the population PRP decreases. But the maximum population PRP is not at zero treatment, since this would correspond to no treatment.

**Figure 10.**
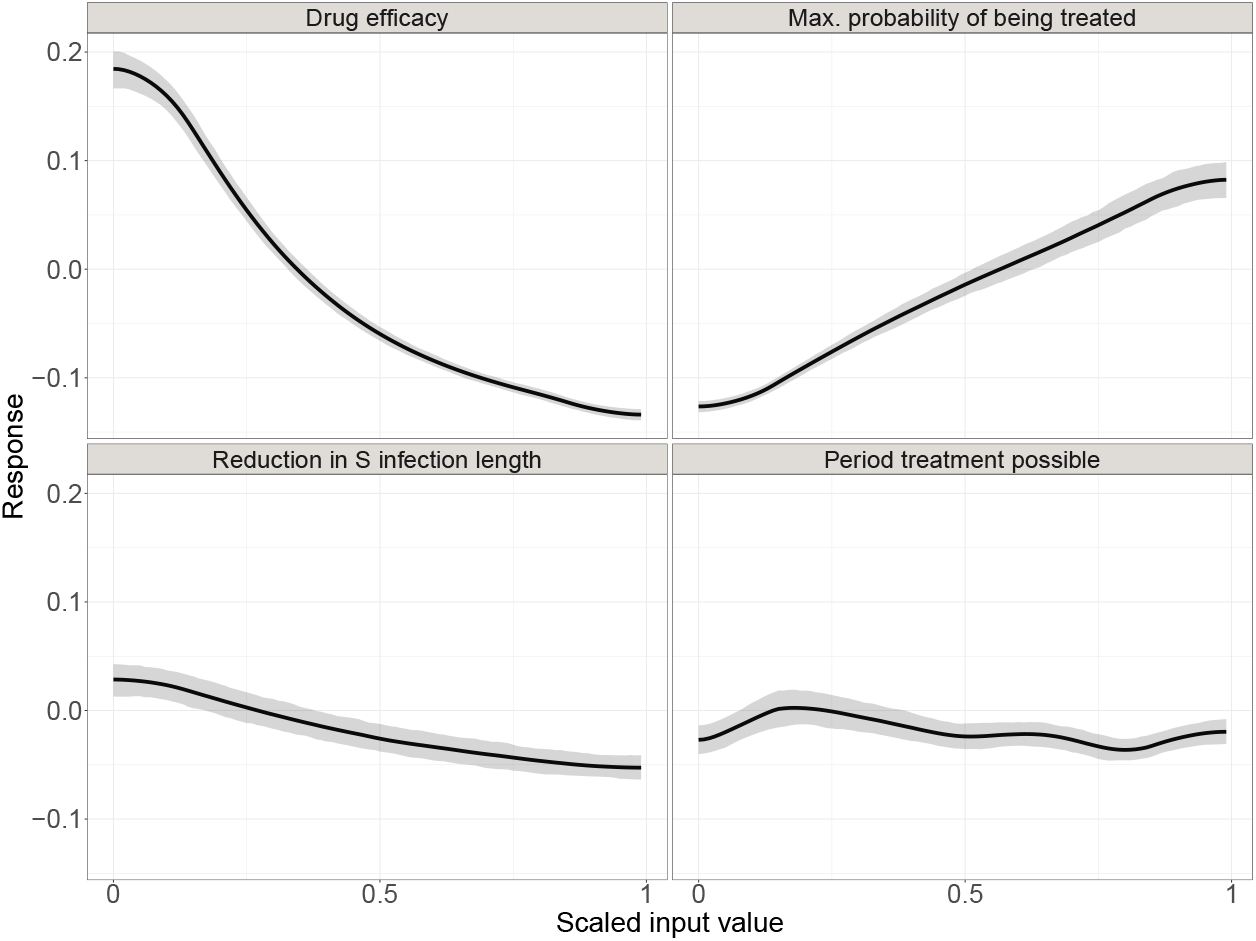
The functional first order effects of the parameters that have the highest sensitivity indices. From Fig 9A, these parameters, shown in the subplot title, are drug efficacy, the maximum probability of being treated, the reduction in sensitive infection length, and the period treatment is possible. The shaded region represents the 90% confidence interval.

Unlike in the case of establishment, where very high and very low drug efficacy was associated with non-establishment, in the case of the spread of resistance the relationship is simpler. The more effective the drug, the less resistance spreads, see Fig 10.

We further compared the competition dynamics with an additional 4000 scenarios in which only the competition dynamics varied. The other parameters were set to the default in Table 1. Of the 4000 scenarios, half are with low transmission (approximately 15% prevalence) and half are with high transmission (approximately 70% prevalence). We conducted a sensitivity analysis on the 1074 (54%) and 1258 (63%) cases where establishment occurred for low and high transmission settings respectively, Fig 11.

**Figure 11.**
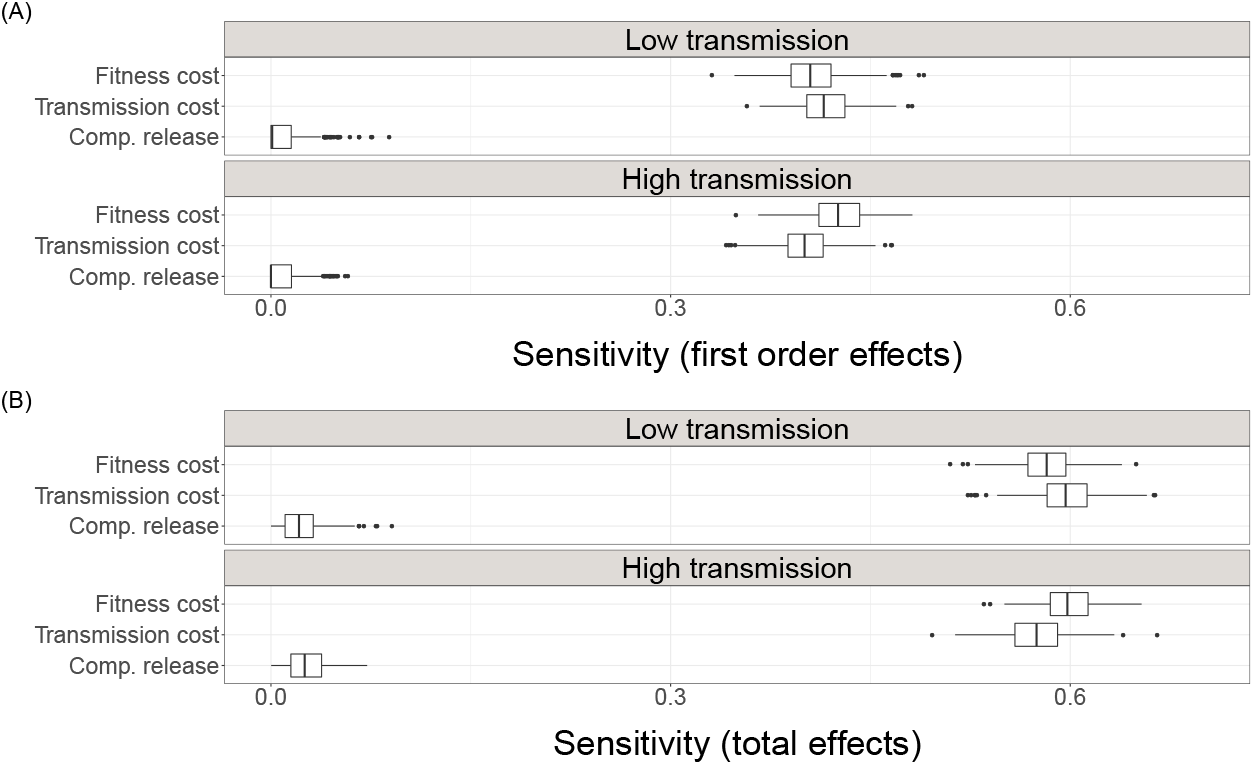
The first order and total sensitivity indices of competition dynamics on the population PRP at 20 years. (A) From 1074 cases where resistance established when the transmission rate was *β* = 0.0004 per day. Median of first order effects sum to 0.826. Median of total effects sum to 1.194. (B) From 1258 cases where resistance established when the transmission rate was *β* = 0.0024 per day. Median of first order effects sum to 0.821. Median of total effects sum to 1.194.

With a focus on the competition dynamics only, we found that ↓ competitive release has no effect. However, we find in low transmission settings, the ↓ fitness cost has a stronger effect at mitigating the spread of resistance, whereas under high transmission settings ↓ transmission cost has a stronger effect. At a fundamental level, it is expected that the effect of the ↓ transmission cost increases in high transmission settings. Nonetheless, it is worth considering what processes the ↓ transmission cost is capturing. For example, with regards to malaria, the ↓ transmission cost relates to recombination, where a high number of mutations required for resistance is a high transmission cost. Therefore, as the transmission rate increases, mosquitoes are carrying more genotypes, and thus the sexual reproduction of the parasite within the mosquito is less likely to generate resistant infections. Also, it is interesting to note that the ordering of the sensitivity to the costs swapped, meaning that not only ↓ is transmission cost more important in high transmission settings than in low transmission, it is more important than the ↓ fitness cost. Whereas in low transmission settings, ↓ fitness cost is more important than ↓ transmission cost.

## Discussion

Without any competition dynamics, resistance establishes and spreads. This is intuitive since treatment clears the sensitive pathogens, and so that eventually only resistant pathogens remain. However this process can be inhibited, or even prevented all together. We presented a novel transmission model to determine what factors prevent establishment, and what factors inhibit the spread. The model tracks the proportion of resistant pathogen, and thus allows for continuous processes, such as varying fitness costs between sensitive and resistant pathogens. From the 12 parameters that varied for the simulations, only values for the transmission rate, the maximum infection length, and the maximum probability of being treated were specific to malaria.

To represent resistant pathogens using the resources which were previously used by the sensitive pathogens, ↑ competitive release, the model linearly increased the proportion of resistant pathogen within a treated host. The primary result of our model is that understanding ↑ competitive release is a priority in preventing resistance establishing in a population. For example, were we reasonable to assume a linear increase of the proportion of resistant pathogen within treated hosts? This is akin to assuming that the selection window [27] occurs immediately after treatment. We conjecture that assuming a sublinear increase, such that shortly after treatment the resistant pathogen ‘replace’ the sensitive pathogen slowly, lessens the effect of ↑ competitive release on resistance establishing. This may be more representative of the selection window occurring some time after initial treatment. Although ↑ competitive release dominates whether resistance establishes, interestingly we showed that it has no impact on the spread of resistance.

The second key result is that, unsurprisingly, treatment drives both the establishment and the spread of resistance. In regards to preventing establishment of resistance, and once established, limiting the spread, we find that only treatment with very high initial efficacy should be used. We expect these results to hold for AMR in general since ↑ competitive release and drug efficacy are general parameters that were not parameterised to malaria specifically. Thirdly, resistance is less likely to establish when infections are short, especially when treatment greatly reduces the length of resistant infections. Then, given establishment, resistance is less likely to spread when treatment greatly reduces the length of sensitive infections. Again, we expect this result to hold for AMR in general. Although the infection length was parameterised to malaria, the comparison of short to long infection lengths still holds, irrespective of the specific boundaries.

We find that although ↑ competitive release means that establishment of resistance is very likely, it is not certain. The factors which make it less likely include a high ↓ transmission cost and a high ↓ fitness cost.

In the Introduction we described two motifs of our model. The first is that we obtain a high-level overview, but with reduced dependence on unknown assumptions. To achieve this, complex mechanisms were included by using simplified functions to capture the broad strokes, so we did not delve into intricacies that would have required many further assumptions. This motif is especially relevant for the competition dynamics (↓ fitness cost, ↓ transmission cost and ↑ competitive release), where each of these are so complex that they warrant a model for each dynamic, likely specific to the infection in question. So instead of these dynamics being emergent properties, we can assign a variable to the competition dynamics directly and vary the strength between extremes, including ‘turning off’ the dynamic. This made a sensitivity analysis into these dynamics straightforward.

The second motif is that we did not model the absolute number of infections, and hence the pathogen load. Instead we modelled the proportion of the pathogen population which is resistant. Hence, the model presented here captures a wider range of mixed infection classes and avoids biases, compared to current compartmental co-infection models. This may be over simplifying the transmission dynamics in cases where the between host transmission is known to be primarily dependent on the pathogen load, and at the same time, the pathogen load within hosts varies greatly across the population. To account for different pathogen loads in different settings, the discretisation of the model could be interpreted as the maximum number of infections that humans can carry, where one assumes that all infected humans carry the maximum. Under this interpretation, throughout this paper we assumed that infected humans carry 10 infections. When interpreting the discretisation as a proxy for pathogen load, it could be argued that the transmission rate and the discretiation should be connected so that as the transmission rate decreases, the discretisation decreases too. This connection adds complexity without providing further insight because the transmission rate is already an approximation. For simplicity, and consistent comparisons, we maintained a standard discretisation throughout.

The ↓ transmission cost encompasses all aspects which make onward transmission of resistant pathogen less likely. We already discussed for malaria, recombination of the parasite in the mosquito is a transmission cost. Furthermore, it may also encompass infection diversity. In high transmission settings there may be more infection diversity, meaning that a person carrying many malaria infections is likely carrying different variants of the parasite. Due to this variety, recombination is more likely to occur, and thus further strengthen the ↓ transmission cost. Since the ↓ transmission cost is already a proxy, separating the cost due to resistance, and the cost due to infection diversity is meaningless. Having discussed the implications of the ↓ transmission cost being a proxy measure, let us similarly discuss the implications for ↑ competitive release and ↓ fitness cost.

As with transmission cost and ↑ competitive release, the ↓ fitness cost captures the main features, but is an approximation. Here we have captured that the ↓ fitness cost comes in the form that, within untreated hosts, (i) the sensitive pathogen population suppresses the resistance, and (ii) resistant pathogens are cleared from the hosts quicker than sensitive pathogens. In this model these two dynamics are captured by one variable *c*. However, perhaps it would be more suitable to model for the *c* in Eq. 8 to be unrelated to Eq. 7. This would capture the two effects of the fitness costs (i) and (ii) separately.

Bushman *et al.* [45] allude to a ↓ fitness cost, whilst excluding multiple infections. The authors mention that when there are many hosts infected with sensitive infections, resistant infections are less able to find an available host, and they state this as a fitness cost. This competition for hosts is excluded here, since at each time step our model infects a proportion of the population with a distribution of resistant pathogen population ***X***_*I*_ (*t*). The obvious next step to the model would be to make it stochastic such that at each time individuals are infected with a given *x_n_* that is randomly drawn from the distribution ***X***_*I*_ (*t*). This would have implications in low transmission areas, where extinction of the resistant pathogen may occur due to stochasticity. However, modelling multiple transmission events with differential equations would greatly increase the model complexity, and thus the model would be better represented as an agent based model.

## Conclusion

We presented a model which includes multiple factors in a general manner, thus making it general to different infections, whilst also encompassing a range of different processes which are known to influence the competition between sensitive and resistant pathogens at the within host and between host level. By tracking the proportion of resistant pathogen we can allow for a finer discretisation of mixed infections than previous models which are either super-infection models, or co-infection models with a 50/50 mix only. This approach is particularly suited to infection where transmission does not depend on pathogen load. Consequently an appropriate infection to explore next with this model would be HIV.

This new modelling approach provided fresh comparisons into which factors are important, and under what settings. We separated our analysis into factors which affect resistance establishment and then, given establishment, factors which affect resistance spread. Our parameterisation focused on malaria, and we reproduce patterns which we know to be true. Namely that ↓ transmission costs have a greater influence in high transmission areas, treatment drives resistance, and to mitigate both the establishment and spread of resistance, treatment should be very effective, where we define efficacy as the proportion of sensitive pathogen cleared upon initial treatment. This last fact is especially relevant for current aims to develop single dose cures of malaria that intend to increase patient adherence [39]. Our model suggests that if single dose cures compromise the effect of the initial clearance of sensitive pathogen, resistance is more likely to establish and spread.

The further insights provided by the model stress that understanding competitive release is imperative in understanding resistance. Although there are many factors which inhibit resistance, as long as resistance thrives within treated hosts, resistance will establish. We need to understand the processes within treated hosts so that, ultimately, drug development and treatment strategy support effective treatment but not resistance. This finding is in agreement with other population models which include resistance [40–42].

## Supporting information

Codes

## Acknowledgments

This research was funded by Melissa Penny’s Swiss National Science Foundation Professorship PP00P3_170702. Calculations were performed at sciCORE (http://scicore.unibas.ch/) scientific computing center at University of Basel.

