## Supplementary material for "The competition dynamics of resistant and sensitive infections": Codes: getCHI.r

```
# Gets distribution of function at each time, for length of infection. 
# Assuming a uniform distribution at the beginning.
# In the model, this uniform distribution is replaced with the relevant distribution.

getCHI <- function(AA, PROP){
  
#  nn                <- 100                          # Discretisation of S to R
  x                 <- seq(0,1,1/nn)                # Proportion which are R
 
  CHI            <- matrix(nrow = nn + 1, ncol = (AA + 1)) 
 
  for (k2 in 1:(AA + 1)){
    res          <- hist(PROP[,k2], breaks = seq(0,1,length = nn + 2), plot = FALSE) 
    CHI[,k2]     <- rev(res$counts / (nn + 1))
  }
  
  return(CHI)
}
  
# NOTE that breaks at c(-diffx, x + diffx) IS WRONG
# for n = 2, difference between c(-0.5, 0.5, 1, 1.5) and 
# c(-x[2], x) = c(-0.5, 0, 0.5, 1) - also wrong. Nothing in first bin.
```
