## Supplementary material for "The competition dynamics of resistant and sensitive infections": Codes: getPROPI_v2.R

```
# The proportion of parasitaemia which are resistant. 
# Changes in time (up to maximum infection length AA) due to competition (cc)

getPROPI_v2 <- function(AA, cc){
  
 # nn                <- 100                          # Discretisation of S to R
  x                 <- seq(0,1,1/nn)                # Proportion which are R
  ell               <- 14                           # Latency period in mosquitoes
  t                 <- seq(0,AA,1)                  # Time vector for infection duration
  Sample            <- FALSE                        # Whether infection
   
  
  ##########################################################
  
  # Proprotion which are resistant - a function that changes in time
  
  ##########################################################
  
  # gradient of proportion at t = 0 is parameterised around 
  # (i) a 50/50 infection having S eliminated after 12ish days. 
  # (ii) a proportion of all resistant (x = 1) is always resistant (gradient = 0)
  # See de Roode, Helinski, Anwar and Read AmNat 2005

  gradient                        <- cc * x - cc
  PROP_R                          <- matrix(nrow = nn + 1, ncol = (AA + 1))
  storehere                       <- nn + 1
  
    for (k1 in 1:(nn + 1)){
    prop_R                      <- gradient[k1] * tt^2 + x[k1]
    prop_R[which(prop_R < 0)]   <- 0
    prop_R[which(prop_R > 1)]   <- 1
    PROP_R[storehere, ]         <- prop_R
    storehere                   <- storehere - 1
  }
  
  rownames(PROP_R)              <- rev(seq(0, 1, 1/nn))
  colnames(PROP_R)              <- seq(0, AA)
  
  # 7 May 2019. Make untreated resistant infections clear quicker
  AR                              <- (1- cc) * AA # infection length of resistant, untreated infections
  AX                              <- round(x * AR + (1 - x) * AA)
  
  PROP_I_v2                       <- matrix(NA, nrow = nn + 1, ncol = (AA + 1))
  here                            <- nn + 1
  for (kx in 1:(nn + 1)){
    PROP_I_v2[here, 1:AX[kx]]   <- PROP_R[here, 1:AX[kx]]
    here <- here - 1
  }
  
  return(PROP_I_v2)
  
}
```
