## Supplementary material for "The competition dynamics of resistant and sensitive infections": Codes: getPROPT_v2.R

```
# Make replacement occur slowly over time. 
# So assume no replacement, and add an increment at each time. 
# If this increment is 0, no replacement, if increment is 1, full replacement. 
# Full replacement means that all cleared S are replaced by R. 
# NOT all S are replaced with R. Some S may remain due to imperfect efficacy


getPROPT_v2 <- function(AA, cc, epsilon, epsilonvec, kappa, Time2RecoverE1, PROP_R_1){
  
 # nn                <- 10                          # Discretisation of S to R
  x                 <- seq(0,1,1/nn)                # Proportion which are R
  
  # Adding half life of drug # v13 addition
  epsilonvec[which(epsilonvec < 0.000000001)] <- 0
  epsilonmat                                  <- t(replicate(nn + 1, epsilonvec))
  
  # Initially epsilon * (1-x) are cleared. That is, 'the epislon proportion of the sensitive proportion'.
  PROP_T_all_top0                             <- PROP_R_1 / ((1 - epsilonmat) * (1 - PROP_R_1) + PROP_R_1) 
  FullReplacementVal                          <- epsilon + PROP_R_1[,1] * (1 - epsilon) 
  #  InitialVal         <- PROP_T_all_top0[,1]
  #  LinearReplacement  <- (FullReplacementVal / InitialVal) * rev(x) + InitialVal
  
  # The infection length is reduced by the proportion which are sensitive to infected - this is recovery. 
  # This matrix will have NA for proportion recovered.
  PROP_T_top0                                 <- matrix(NA, nrow = nn + 1, ncol = (AA + 1))
  here                                        <- nn + 1
  for (kx in 1:(nn + 1)){
    PROP_T_top0[here, 1:Time2RecoverE1[kx]]   <- PROP_T_all_top0[here, 1:Time2RecoverE1[kx]]
    here <- here - 1
  }
  
  # Linearly (in time) shifting proportion towards higher x due to replacement
  PROP_T_top                                  <- PROP_T_top0
  for (kt in 2:(AA+1)){
    PROP_T_top[, kt]                          <- PROP_T_top0[,(kt-1)] + kappa
  }
  
  # The shift up cannot be more than 'full replacement' proportion
  for (kx in 1:(nn + 1)){
    idx                                       <- which(PROP_T_top[kx,] >  FullReplacementVal[kx])
    PROP_T_top[kx, idx]                       <- FullReplacementVal[kx]
  }
  
  return(PROP_T_top)
}

#  # When given treatment, 'epsilon' of the Sensitive proportion is cleared, and the rest is resistant.
#PROP_T_all_top0  <- replicate(AA, rev(x)) / ((1 - epsilon) * (1 - replicate(AA, rev(x))) + replicate(AA, rev(x))) #epsilon + PROP_R_1 * (1 - epsilon)
#PROP_T_all_top1  <- epsilon + replicate(AA, rev(x)) * (1 - epsilon) #epsilon + PROP_R_1 * (1 - epsilon)
```
