## Supplementary material for "The competition dynamics of resistant and sensitive infections": Codes: getrvec.r

```
getrvec <- function(AA, rx, DaysTreatmentPoss){
  
  tt                        <- seq(0, AA, 1)
  t_hat                     <- pi/(DaysTreatmentPoss) * tt[1:DaysTreatmentPoss]
  rvec                      <- matrix(0, nrow = (AA + 2), ncol = 1) 
  rvec[1:DaysTreatmentPoss] <- rx * sin(t_hat + sin(t_hat))
  
  return(rvec)
}
```
