## Supplementary material for "The competition dynamics of resistant and sensitive infections": Codes: getShiftVec.R

```
# A vector which shifts an infection(s) with resistant parasitaemia x, towards a higher x due to treatment clearing S

getShiftVec <- function(NewProp){
  
 # nn                <- 100                          # Discretisation of S to R
  x                 <- seq(0,1,1/nn)                # Proportion which are R
  Sample            <- FALSE                        # Whether infection
  
  ShiftVec         <- matrix(nrow = nn + 1, ncol = 1) 
  
  for (k1 in 1:(nn + 1)){
    res            <- hist(NewProp[k1], breaks = seq(0,1,length = nn + 2), plot = FALSE) 
    ShiftVec[k1]   <- which(rev(res$counts) == 1)
  }
  
  return(ShiftVec)
}
```
