## Supplementary material for "The competition dynamics of resistant and sensitive infections": Codes: getTime2Recover.r

```
getTime2Recover <- function(AA, rhoR, rhoS, epsilon){
  
  #nn               <- 100                          # Discretisation of S to R
  x                <- seq(0, 1, 1/nn) # proportion which are Resistant
  
  # Reduced time of infection for treated hosts
  denomR           <- 1 + rhoR * epsilon^2
  denomS           <- 1 + rhoS * epsilon^2
  
  Time2RecoverE1   <- round(x * (AA/denomR) + (1 - epsilon) * (1 - x) * (AA/denomS))
  
  return(Time2RecoverE1)
}
```
