## Supplementary material for "The competition dynamics of resistant and sensitive infections": Codes: Supplementary_NotAsFunction.R

```
# Can turn off sampling. Quicker. 
# Add treatment after Steady State
# ============================================================================================


library(ggplot2)
library(reshape2)
library(gridExtra)
library(purrr)


source("getPROPI_v2.R")
source("getPROPT_v2.R")
source("getrvec.R")
source("getTime2Recover.R")
source("getCHI.R")
source("getShiftVec.R") 

#cc                <- runif(1, 0, 1)                # Competition [0 = none, 1 = max] 
#kappa             <- runif(1, 0, 1)                # Replacement [0 = none, 1 = max]
#recombo           <- runif(1, 0, 1)                # Recombination [0 = none OVERALL = TRANSMIT, 1 = max, TRANSMIT no R]
#beta              <- runif(1, 0.0001, 0.01)        # Transmission rate [0.0001, 0.01]
#PopSize           <- 10000                         # Only relevant if Sample = TRUE
#AA                <- sample(50:250,1)              # Length of untreated infection [50, 250]
#rx                <- runif(1, 0.0039, 0.11)        # Max. probability of being treated [0, 1]
#DaysTreatmentPoss <- sample(1:AA,1)            # Days treatment possible [1, AA]
#inv_alpha         <- DaysTreatmentPoss/AA          # Proportion of infection time that treatment poss
#epsilon           <- runif(1, 0, 1)                # Drug efficacy [0,1]
#rhoR              <- runif(1, 0, 1)                # Reduction in infn length of R [0, 1]
#rhoS              <- sample(1:19,1)                # Reduction in infn length of S [1, 10]
#lambda1           <- runif(1, 0, 1)                # Drug half-life [0, 1]
#lambda2           <- runif(1, 1, 10)               # Drug adherence [1 = none, 10] 
#input              <- read.table("output_v31a_changeDynamics_beta0001_rx07.csv", sep = ";", header = TRUE)
cc                 <- 0#.004#input$cc[ks]#0.004#$cc[ks]
kappa              <- 0#0#input$kappa[ks]
recombo            <- 0#input$recombo[ks]
beta               <- 0.0002#input$beta[ks]
PopSize            <- 10000#as.numeric(as.character(input$PopSize[ks]))
AA                 <- 140#as.numeric(as.character(input$AA[ks]))
rx                 <- 0.24#0.01#input$rx[ks]
inv_alpha          <- 1/14#as.numeric(as.character(input$inv_alpha[ks]))
DaysTreatmentPoss  <- round(inv_alpha * AA)
epsilon            <- 0.9#as.numeric(as.character(input$epsilon[ks]))
rhoR               <- 0.2#as.numeric(as.character(input$rhoR[ks]))
rhoS               <- 2#as.numeric(as.character(input$rhoS[ks]))
lambda1            <- 0.4#as.numeric(as.character(input$lambda1[ks]))
lambda2            <- 4#as.numeric(as.character(input$lambda2[ks]))

##########################################################

# Model parameters

##########################################################
NYrs              <- 30
NDays             <- 365 * NYrs
SStime            <- 365 * 3

nn                <- 10                          # Discretisation of S to R
x                 <- seq(0,1,1/nn)                # Proportion which are R
ell               <- 14                           # Latency period in mosquitoes
tt                <- seq(0, AA, 1)                # Time vector for infection duration

Sample            <- FALSE                        # Whether infected

##########################################################

# Set - up

##########################################################

# Drug efficacy changing in time - Weibull decay https://github.com/SwissTPH/openmalaria/wiki/ModelDecayFunctions
epsilonvec        <- epsilon * exp(-(lambda1 * tt)^lambda2 * log(2)) 

Time2RecoverE1    <- getTime2Recover(AA, rhoR, rhoS, epsilon)

# Probability of getting treated 
rvec              <- getrvec(AA, rx, DaysTreatmentPoss)
not_rvec          <- 1 - rvec

# Proportion of resistant infections - changes for length of infection.
PROP_R            <- getPROPI_v2(AA, cc)
PROP_T_top        <- getPROPT_v2(AA, cc, epsilon, epsilonvec, kappa, Time2RecoverE1,   PROP_R) 

# Distribution of Resistant proportion of infections - changes in time
CHI_I             <- getCHI(AA, PROP_R)
CHI_T             <- getCHI(AA, PROP_T_top)

##########################################################

# The probability of recovering at time a, when not given treatment

##########################################################

mmatrixI                   <- PROP_R
mmatrixI[!is.na(mmatrixI)] <- 1
mmatrixI[is.na(mmatrixI)]  <- 0 

##########################################################

# The probability of recovering at time a, when given treatment

##########################################################

ShiftVec                   <- getShiftVec(PROP_T_top) # The shift due to treatment
mmatrixT                   <- PROP_T_top
mmatrixT[!is.na(mmatrixT)] <- 1
mmatrixT[is.na(mmatrixT)]  <- 0 


##########################################################

# Initial number of hosts in each category

##########################################################

Susceptible           <- matrix(PopSize, nrow = 1, ncol = NDays)
Infected              <- matrix(0, nrow = 1, ncol = NDays)
Infected[1:ell]       <- 2/100 * PopSize                       # 2 percent infected 
Susceptible[1:ell]    <- PopSize - Infected[1]
Treated               <- matrix(0, nrow = 1, ncol = NDays)
NewInfn               <- Susceptible * beta 
NewTreated            <- matrix(0, nrow = NDays, ncol = NDays)

Resistant             <- matrix(NA, nrow = 1, ncol = NDays)
Sensitive             <- matrix(NA, nrow = 1, ncol = NDays)

##########################################################

# Initial distribution of proportion of resistant infections 

##########################################################

OVERALLDIST                 <- matrix(0, nrow = (nn + 1), ncol = NDays)
OverallInit                 <- matrix(0, nrow = (nn + 1), ncol = 1)
OverallInit[1]              <- 0.0001 # initial resistance
OverallInit[nn + 1]         <- 1 - OverallInit[1]
OVERALLDIST[, 1:SStime]     <- OverallInit
TRANSMITDIST                <- OVERALLDIST

INFECTED_DIST               <- matrix(0, nrow = (nn + 1), ncol = NDays)
TREATED_DIST                <- matrix(0, nrow = (nn + 1), ncol = NDays)
INFECTED_DIST[, 1:SStime]   <- OverallInit # begin with little resistance
TREATED_DIST[, 1:SStime]    <- OverallInit # begin with little resistance

RandomInfecteds             <- matrix(0, nrow = (nn + 1), ncol = NDays)  
RandomInfecteds[, 1:SStime] <- OverallInit

ShiftedTdist                <- array(rep(NaN, (NDays * NDays * (nn+1))), dim=c(NDays, NDays, (nn+1))) # x for the individuals infected
PluckedTdist                <- array(rep(NaN, (NDays * NDays * (nn+1))), dim=c(NDays, NDays, (nn+1))) # The shifted x after to treatment

SteadyState                 <- array(1, dim = c(1, NDays))

DF_PROP_T                   <- data.frame(PROP_T_top)
colnames(DF_PROP_T)         <- seq(0,AA)
meltDF_PROP_T               <- melt(t(DF_PROP_T), id = rownames(PROP_T_top))

EpsilonDF                   <- data.frame(cbind(seq(0, AA), epsilonvec))
colnames(EpsilonDF)         <- c("Var1", "Efficacy")

##########################################################

# No resistance until Steady State

##########################################################

for (kt in 1:ell){
  # Need to store initial conditions here for later
  PluckedTdist[kt, 1:SStime, ] <- matrix(OverallInit, nrow=SStime, ncol=length(OverallInit), byrow=TRUE)
  ShiftedTdist[kt, 1:SStime, ] <- matrix(OverallInit, nrow=SStime, ncol=length(OverallInit), byrow=TRUE)
  
}

##########################################################

# And go!! 

##########################################################

for (kt in (ell + 1):NDays){
  #  kt     <- ell
  #  while (abs(SteadyState[kt]) > 0) { 
  #      kt <- kt + 1
  #     if (kt > NDays){break}
  if (kt %% 100 == 0){print(kt)}
  
  start                     <- max(ell, (kt - AA))
  end                       <- kt - 1
  
  Infected_dist             <- matrix(0, nrow = (nn + 1), ncol = 1)
  
  counter                   <- 1
  
  NewInfn[kt]               <- Susceptible[kt - 1] * beta  # bites at each time
  
  RandomInfecteds[ ,kt]     <- TRANSMITDIST[, kt - ell]
  
  for (kq in start:end){
    
    DaysSinceInfn           <- kt - kq
    
    NotRecoveredI           <- NewInfn[kq] * sum(TRANSMITDIST[, (kq - ell + 1)] * mmatrixI[, DaysSinceInfn])
    
    Infected[kt]            <- Infected[kt] + NewInfn[kq] * prod(not_rvec[1:(counter + 1)]) * NotRecoveredI
    # Number of people infected on day kq, treated on day kt
    # Infected_kq_DaysAgo <- NewInfn[kq] * prod(not_rvec[1:DaysSinceInfn]) * rvec[(DaysSinceInfn + 1)]
    NewTreated[kq, kt]      <- NewInfn[kq] * prod(not_rvec[1:DaysSinceInfn]) * rvec[(DaysSinceInfn + 1)]#
    
    Infected_dist           <- Infected_dist +                  # sum of ...
      NewInfn[kq] / Infected[kt]  *    # ratio of those infected on day kq to total infected today [kt]... 
      prod(not_rvec[1:counter]) *      # that have not been treated ...
      RandomInfecteds[ , kt] *         # had distn from TRANSMITDSIT[kq] ...
      NotRecoveredI *                  # not recovered ... 
      CHI_I[, kt - kq]                 # and competition has altered distn since kq
    
    if (sum(Infected_dist, na.rm = T) > 0){Infected_dist <- (Infected_dist) / sum(Infected_dist)}
    
    PluckedFrom              <- TRANSMITDIST[, (kq - ell + 1)] * CHI_I[, DaysSinceInfn]#counter]
    PluckedFrom              <- PluckedFrom / sum(PluckedFrom)
    PluckedT                 <- PluckedFrom # is a vector (matrix in first 'if' part) [R -> S]
    ShiftedT                 <- matrix(0, nrow = (nn + 1), ncol = 1)
    idx                      <- which(PluckedT > 0)
    ShiftedT[ShiftVec[idx,]] <- PluckedT[idx] # is a vector (matrix in first 'if' part)
    # Distn of those infected on day kq, treated on day kt (before and after treatment)
    PluckedTdist[kq, kt, ]   <- PluckedT / sum(PluckedT) # distribution at time of treatement
    ShiftedTdist[kq, kt, ]   <- ShiftedT / sum(ShiftedT) # should already sum to 1
    
    Treated_dist             <- matrix(0, nrow = (nn + 1), ncol = 1)
    
    if ((NewInfn[kq]) > 0.5){
      
      for (kk in 0:(kt - kq - 1)){
        
        # infected on day kq, treated on day kq + kk + 1
        #idx                    <- kk + 1
        #scaling_factor         <- (kt - kq - kk - 1) / AA
        #mmatrixidx             <- min(AA, round(idx * scaling_factor))
        mmatrixTidx             <- min(AA, round((AA * (kt - kq - kk + 1)) / (AA - kk)))
        
        NotRecoveredT          <- NewTreated[kq, kq + kk + 1] * sum(PluckedTdist[kq, kq + kk + 1, ] * mmatrixT[, mmatrixTidx]) 
        
        AddingTreated          <- NewInfn[kq] *                 # Those infected on day kq ..
          prod(not_rvec[1:(kk+1)]) *    # .. who were not treated for kk days, 
          rvec[kk+2] *                  # .. and then treated
          NotRecoveredT                 # .. and not recovered since day kq + kk + 1
        
        Treated[kt]            <- Treated[kt]  + AddingTreated # sum of above
        
        if (Treated[kt] > 0 )
        {
          Treated_dist         <- Treated_dist +                      # Sum of ...
            AddingTreated / Treated[kt] *       # ... proportion of those infected on day kq and treated on day kq + kk + 1
            NotRecoveredT *                      # ... those not recovered...
            ShiftedTdist[kq, kq + kk + 1, ] *   # ... with shifted distribution...
            CHI_T[, mmatrixTidx]                 # ... and competition and replacement has altered distn
          if (sum(Treated_dist, na.rm = T) > 0) {Treated_dist <- (Treated_dist) / sum(Treated_dist)
          } else {
            Treated_dist          <- matrix(0, nrow = (nn + 1), ncol = 1)
          } }
      } # close kk for
    } # close if
  } # close kq for
  
  Susceptible[kt]               <- PopSize - Infected[kt] - Treated[kt]
  
  if (kt > SStime){
    #=================================================================================
    # Multiple bites - don't change infection length, but shift towards OVERALL by factor beta
    
    Infected_dist                 <- (TRANSMITDIST[,kt - ell] * beta) + (Infected_dist * (1 - beta))
    Treated_dist                  <- (TRANSMITDIST[,kt - ell] * beta) + (Treated_dist * (1 - beta))
    
    #=================================================================================
  } else {
    Infected_dist <- matrix(0, nrow = 1, ncol = NDays)
    Treated_dist  <- matrix(0, nrow = 1, ncol = NDays)
  }
  
  if (sum(Infected_dist, na.rm = T) > 0){INFECTED_DIST[,kt]  <- Infected_dist/sum(Infected_dist)}
  if (sum(Treated_dist, na.rm = T)  > 0){TREATED_DIST[,kt]   <- Treated_dist/sum(Treated_dist)}
  
  OVERALLDIST[, kt]             <- Infected[kt] * INFECTED_DIST[,kt] + Treated[kt] * TREATED_DIST[,kt] 
  OVERALLDIST[which(OVERALLDIST[, kt] < 0), kt]        <- 0 
  if (sum(OVERALLDIST[, kt]) > 0){OVERALLDIST[, kt]    <- OVERALLDIST[, kt] / sum(OVERALLDIST[, kt])}
  
  Resistant[kt]                 <- sum(rev(x) * OVERALLDIST[,kt])
  Resistant[kt]                 <- round(Resistant[kt],4)
  Sensitive[kt]                 <- sum((1-rev(x)) * OVERALLDIST[,kt])
  if (Resistant[kt] > OverallInit[1]){
    SteadyState[kt]               <- mean(c(Resistant[kt] - Resistant[(kt-10):(kt-1)]), na.rm = TRUE)  
  }
  
  # Recombination - applied to each 'bar' or 'density for a given x'
  #TRANSMITDIST[, kt]           <- ((1 - recombo) * rev(x) * OVERALLDIST[, kt]) +  ((1 - rev(x)) * OVERALLDIST[, kt])
  TRANSMITDIST[, kt]            <- ((1 - recombo) * OVERALLDIST[, kt]) 
  TRANSMITDIST[(nn + 1), kt]    <- 1 - sum(TRANSMITDIST[(1:nn), kt])
  
} # end day kt loop

#if (kt > NDays){
#  SteadyStateTime <- kt
#} else{
#  SteadyStateTime <- min(which(SteadyState[1,] == 0)) - SStime
#}
#establishTime   <- min(which(Resistant[SStime:NDays] > OverallInit[1]))
RR <- Resistant[1,]
Est <- which(RR == max(RR, na.rm = T))
SteadyStateTime <- min(Est)

NotEst <- which(RR == min(RR, na.rm = T))
establishTime   <- max(NotEst)
  # ============================================================================================

DF                                <- data.frame(cbind(rev(x), 
                                     INFECTED_DIST[,NDays], 
                                     TREATED_DIST[,NDays], 
                                     OVERALLDIST[,NDays], 
                                     TRANSMITDIST[,NDays]))
colnames(DF) <- c("x", "Infected", "Treated", "Overall", "Transmit")
DFmelt <- melt(DF, id = 'x')
q  <-  ggplot(data = DFmelt, aes(x = x, y = value)) + #, fill = variable)) +
       geom_bar(stat = "identity", position = "dodge", alpha = 1) + theme_bw() +
      #scale_fill_manual(values=c("brown4", "slateblue","orchid4")) +  
      #scale_fill_manual(values=c("lightgray", "lightgray","black")) +  
      # scale_fill_manual(values=c("brown4", "darkolivegreen","black", "black")) +  
       facet_grid(~ variable) +
       theme(strip.text.x = element_text(size=48)) +
       scale_y_continuous(name = "Density", limits = c(0,1)) +
       scale_x_continuous(name = "Resistant prop.", breaks=seq(0,1,0.5) , labels= seq(0,1,0.5)) +
       theme(axis.text.x = element_text(size=36),
       axis.text.y = element_text(size=36)) +
       theme(legend.position = "none") + 
       theme(axis.title = element_text(size=42)) + 
       theme(strip.background = element_rect(fill = alpha('black', 0.2))) +
       theme(strip.text = element_text(colour = 'black')) 
q


DF                                <- data.frame(cbind(rev(x), 
                                                      OVERALLDIST[,NDays], 
                                                      TRANSMITDIST[,NDays]))
colnames(DF) <- c("x", "From humans", "To humans")
DFmelt <- melt(DF, id = 'x')
q  <-  ggplot(data = DFmelt, aes(x = x, y = value)) + #, fill = variable)) +
  geom_bar(stat = "identity", position = "dodge", alpha = 1) + theme_bw() +
  facet_grid(~ variable) +
  theme(strip.text.x = element_text(size=32)) +
  scale_y_continuous(name = "Density", limits = c(0,1)) +
  scale_x_continuous(name = "Resistant prop.", breaks=seq(0,1,0.5) , labels= seq(0,1,0.5)) +
  theme(axis.text.x = element_text(size=36),
        axis.text.y = element_text(size=36)) +
  theme(legend.position = "none") + 
  theme(axis.title = element_text(size=42)) + 
  theme(strip.background = element_rect(fill = alpha('#604c38', 0.2))) +
  theme(strip.text = element_text(colour = 'black')) 
q

#annotate("rect", ymin = -Inf, ymax = Infected[NDays]/(Treated[NDays] + Infected[NDays]), xmin = -Inf, xmax = Inf, fill = 'brown4', alpha = 0.2) + 
#annotate("rect", ymin = Infected[NDays]/(Treated[NDays] + Infected[NDays]), ymax = Inf, xmin = -Inf, xmax = Inf, fill = 'slateblue', alpha = 0.2) + 
  
  
##########################################################
  
DF_PROP_T <- data.frame(PROP_T_top)
colnames(DF_PROP_T)<- seq(0,AA)
meltDF_PROP_T    <-  melt(t(DF_PROP_T), id = rownames(PROP_T_top))
  
EpsilonDF <- data.frame(cbind(seq(0,AA), epsilonvec))
colnames(EpsilonDF)<- c("Var1", "Efficacy")
  
ggplot(data = meltDF_PROP_T, aes(x = Var1, y = value, group = Var2, colour = Var2)) + 
  geom_line(size = 2) +
  geom_line(data = EpsilonDF, aes(x = Var1, y = Efficacy), inherit.aes = FALSE, size = 2, linetype = "dashed", colour = "darkgreen" ) + #https://stackoverflow.com/questions/44988682/scatterplot-error-in-funxi-object-group-not-found
  #annotate("rect", ymin = -Inf, ymax = Inf, xmin = -Inf, xmax = Inf, fill = 'darkolivegreen', alpha = 0.2) + 
  scale_y_continuous(limits = c(0,1), name = "Resistant prop.", breaks = seq(0,1,0.5)) + theme_bw() +
  scale_x_continuous(name="Infection time", breaks=c(0,AA) , labels= c("A - T", "A") ) +
  theme(axis.text.x = element_text(size=32),
        axis.text.y = element_text(size=32)) +
  theme(legend.position="none") + 
  theme(axis.title=element_text(size=32)) 

DF_PROP_1        <- data.frame(PROP_R)
colnames(DF_PROP_1)<- seq(0,AA)
meltDF_PROP_1    <- melt(t(DF_PROP_1), id = rownames(PROP_R))
ggplot(meltDF_PROP_1 , aes(x = Var1, y = value, group = Var2, colour = Var2)) + 
  geom_line(size = 2) +
 # annotate("rect", ymin = -Inf, ymax = Inf, xmin = -Inf, xmax = Inf, fill = 'brown4', alpha = 0.2) + 
  scale_y_continuous(limits = c(0,1), name = "Resistant prop.", breaks = seq(0,1,0.5))  + theme_bw() +
  scale_x_continuous(name="Infection time", breaks=c(0,AA) , labels= c("0", "A") ) +
  theme(axis.text.x = element_text(size=32),
        axis.text.y = element_text(size=32)) +
  theme(legend.position="none") + 
  theme(axis.title=element_text(size=32))

MeanResistance <- matrix(NA, nrow = NDays, ncol = 1)
for (k in 1:(NDays - 100)){
  MeanResistance[k + 100] <- mean(Resistant[1,k:(k+100)], na.rm = TRUE)
}
#DF                  <- data.frame(cbind(seq(1,NDays-100), MeanResistance))
DF                  <- data.frame(cbind(seq(1,NDays), Resistant[1,]))
colnames(DF)        <- c("Time", "Resistant")
meltDF              <- melt(DF, id = "Time")
q <-  ggplot(data = meltDF, aes(x = Time, y = value, colour = variable)) +
  geom_line(size = 3) + theme_bw() + 
  theme(axis.title.x = element_text(margin = margin(t = 20))) + 
  theme(axis.title.y = element_text(margin = margin(r = 20))) + 
  scale_colour_manual(values = c("gray", "black")) + 
  scale_x_continuous(name = "Time (years)", breaks = seq(0, NDays, 365*5), labels = seq(0, 30, by = 5)) +
  scale_y_continuous(name = "Resistant prop.") +
  theme(legend.position="none") +
  theme(axis.text.x = element_text(size=60),
        axis.text.y = element_text(size=60)) +
  theme(axis.title = element_text(size=60))  +
  geom_vline(xintercept = SStime, size = 5, alpha = 0.8, linetype = "dashed")
q
#write.table(DF, file = "Model_v33_Resistance_nn5__beta00003_rx008.csv", sep = ";")
#ggsave('Model_v33_Resistance_nn5__beta00003_rx008.png', height = 8, width = 16)
#ggsave('Model_v33_Resistance_nn5__beta00003_rx008.pdf', height = 8, width = 16)
#ggsave('Model_v33_Resistance_nn5__beta00003_rx008.tiff', height = 8, width = 16)
  
  
DF                  <- data.frame(cbind(Infected[ell+1:NDays]/PopSize * 100, Treated[ell+1:NDays]/PopSize * 100, seq(ell + 1,NDays)))
colnames(DF)        <- c("Infected", "Treated", "Day")
meltDF              <-  melt(DF, id = "Day")
pv                  <- ggplot(meltDF, aes(x=Day, y=value, fill=variable)) + geom_area(alpha=0.6) + theme_bw() 
pv <- pv  + scale_fill_manual(values=c("brown4", "slateblue")) + 
      theme(axis.title.x = element_text(margin = margin(t = 20))) + 
      theme(axis.title.y = element_text(margin = margin(r = 20))) + 
      scale_x_continuous(name = "Time (years)",  breaks = seq(0, NDays, 365*5), labels = seq(0, NYrs, by = 5)) +
      scale_y_continuous(name="% of hosts", limits = c(0,2)) +
      #theme(legend.position="none") +
      guides(fill=guide_legend(title="")) + 
      theme(legend.position = c(0.8,0.6)) + #, legend.direction = "horizontal",  legend.spacing.x = unit(1.0, 'cm')) + 
      theme(legend.text=element_text(size=70)) + 
      theme(axis.text.x = element_text(size=60),
            axis.text.y = element_text(size=60)) +
      theme(axis.title = element_text(size=60)) +
      geom_vline(xintercept = SStime, size = 5, alpha = 0.8, linetype = "dashed")
pv
#write.table(DF, file = "Model_v33_Nhosts_nn5__beta00003_rx008.csv", sep = ";")
#ggsave('Model_v33_Nhosts_nn5__beta00003_rx008.png', height = 8, width = 16)
#ggsave('Model_v33_Nhosts_nn5__beta00003_rx008.pdf', height = 8, width = 16)
#ggsave('Model_v33_Nhosts_nn5__beta00003_rx008.tiff', height = 8, width = 16)

DF1                  <- data.frame(cbind(seq(1,NDays), Resistant[1,]))
DF1                  <-  DF1[-(1:ell), ]
DF1 <- cbind(DF1, rep("Resistant prop.", nrow(DF1)))
colnames(DF1) <- c("Day", "value", "variable")

DF2  <- data.frame(cbind(Infected[ell+1:NDays - ell]/PopSize * 100, Treated[ell+1:NDays - ell]/PopSize * 100, seq(ell + 1,NDays - ell)))
colnames(DF2)        <- c("Infected", "Treated", "Day")

DF12 <- cbind(DF1, DF2[-(1:ell), ])

#DF12 <- cbind(DF1[(ell+1:NDays), ], DF2)
MaxY <- 6#max((DF2$Infected + DF2$Treated), na.rm = TRUE) # 6
MaxR <- max(Resistant[1,], na.rm = TRUE)
DF1$value <- DF1$value * (MaxY * 1/MaxR)


meltDF2              <- melt(DF2, id = "Day")
meltDF1              <- melt(DF1, id = "Day")

# Darjeeling1
# orange = "#F98400", blue = "#5BBCD6", red = "#FF0000"

pv                  <- ggplot(meltDF2, aes(x = Day, y = value, fill = variable)) + 
  geom_area( position = 'stack', alpha = 1) + theme_bw() 
pv <- pv  + 
  geom_line(data = DF1, size = 2, color = "#FF0000") + 
 # scale_fill_manual(values=c("#F98400", "#FF0000", "#5BBCD6"), breaks=c("Infected","Treated","Resistant prop.")) + 
  scale_fill_manual(values=c("#F98400", "#5BBCD6", "#FF0000"), breaks=c("Infected","Treated","Resistant prop.")) + 
  theme(axis.title.x = element_text(margin = margin(t = 20))) + 
  theme(axis.title.y = element_text(margin = margin(r = 20))) + 
  labs(tag = "(A)") + #ggsave('Model_v33_NhostsResistance_beta00001_rx024.png', height = 6, width = 18)
  #labs(tag = "(B)") + #ggsave('Model_v33_NhostsResistance_beta00001_rx024_KappaOn.png', height = 6, width = 18)
  #labs(tag = "(C)") + #ggsave('Model_v33_NhostsResistance_beta00001_rx024_DynamicsOn.png', height = 6, width = 18)
  theme(plot.tag = element_text(size = 24)) +
  scale_x_continuous(name = "Time (years)",  breaks = seq(0, NDays, 365*5), labels = seq(0, NYrs, by = 5)) +
  #scale_y_continuous(name="% of hosts",
  #                  sec.axis = sec_axis(~. / (MaxY * 1/MaxR), name = "Resistant prop.", breaks = c(0, MaxR))) +
  scale_y_continuous(name="% of hosts",  limits = c(0,7), breaks = c(0, 3, 6),
                    sec.axis = sec_axis(~.  / (MaxY * 1), name = "Resistant prop.", breaks = c(0, 1))) +
  theme(axis.text.y.right = element_text(color = "#FF0000")) +
  theme(legend.position="none") +
  guides(fill=guide_legend(title="")) + 
  theme(legend.position = c(0.41,0.82), legend.direction = "horizontal",  legend.spacing.x = unit(0.8, 'cm')) + 
  theme(legend.text=element_text(size=28)) + 
  theme(axis.text.x = element_text(size=36),
        axis.text.y = element_text(size=36)) +
  theme(axis.title = element_text(size=36)) +
  geom_vline(xintercept = SStime, size = 5, alpha = 0.8, linetype = "dashed") 
pv

# 29 May 2020

write.table(DF12, file = "Model_v33_Resistance_beta00002_rx024.csv", sep = ";")
ggsave('Model_v33_NhostsResistance_beta00002_rx024.png', height = 6, width = 18)
ggsave('Model_v33_NhostsResistance_beta00002_rx024.pdf', height = 6, width = 18)
ggsave('Model_v33_NhostsResistance_beta00002_rx024.tiff', height = 6, width = 18)

#write.table(DF12, file = "Model_v33_Resistance_beta00002_rx024_KappaOn.csv", sep = ";")
#ggsave('Model_v33_NhostsResistance_beta00002_rx024_KappaOn.png', height = 6, width = 18)
#ggsave('Model_v33_NhostsResistance_beta00002_rx024_KappaOn.pdf', height = 6, width = 18)
#ggsave('Model_v33_NhostsResistance_beta00002_rx024_KappaOn.tiff', height = 6, width = 18)

#write.table(DF12, file = "Model_v33_Resistance_beta00002_rx024_DynamicsOn.csv", sep = ";")
#ggsave('Model_v33_NhostsResistance_beta00002_rx024_DynamicsOn.png', height = 6, width = 18)
#ggsave('Model_v33_NhostsResistance_beta00002_rx024_DynamicsOn.pdf', height = 6, width = 18)
#ggsave('Model_v33_NhostsResistance_beta00002_rx024_DynamicsOn.tiff', height = 6, width = 18)
#==============================================================

#write.table(DF12, file = "Model_v33_Resistance_beta00001_rx024.csv", sep = ";")
#ggsave('Model_v33_NhostsResistance_beta00001_rx024.png', height = 6, width = 18)
#ggsave('Model_v33_NhostsResistance_beta00001_rx024.pdf', height = 6, width = 18)
#ggsave('Model_v33_NhostsResistance_beta00001_rx024.tiff', height = 6, width = 18)

#ggsave('Model_v33_NhostsResistance_beta00001_rx024_DynamicsOn.png', height = 6, width = 18)
#ggsave('Model_v33_NhostsResistance_beta00001_rx024_DynamicsOn.pdf', height = 6, width = 18)
#ggsave('Model_v33_NhostsResistance_beta00001_rx024_DynamicsOn.tiff', height = 6, width = 18)


#write.table(DF, file = "Model_v33_Resistance_nn5__beta00003_rx008.csv", sep = ";")
#ggsave('Model_v33_NhostsResistance_nn5__beta00003_rx008.png', height = 8, width = 16)
#ggsave('Model_v33_NhostsResistance_nn5__beta00003_rx008.pdf', height = 8, width = 16)
#ggsave('Model_v33_NhostsResistance_nn5__beta00003_rx008.tiff', height = 8, width = 16)
```
